# Structure and Dynamic Basis of Molecular Recognition Between Acyltransferase and Carrier Protein in *E. coli* Fatty Acid Synthesis

**DOI:** 10.1101/2020.05.15.098798

**Authors:** Laetitia E. Misson, Jeffrey T. Mindrebo, Tony D. Davis, Ashay Patel, J. Andrew McCammon, Joseph P. Noel, Michael D. Burkart

## Abstract

Fatty acid synthases (FASs) and polyketide synthases (PKSs) iteratively elongate and often reduce two-carbon ketide units in *de novo* fatty acid and polyketide biosynthesis. Cycles of chain extensions in FAS and PKS are initiated by an acyltransferase (AT), which loads monomer units onto acyl carrier proteins (ACPs), small, flexible proteins that shuttle covalently linked intermediates between catalytic partners. Formation of productive ACP-AT interactions is required for catalysis and specificity within primary and secondary FAS and PKS pathways. Here, we use the *Escherichia coli* FAS AT, FabD, and its cognate ACP, AcpP, to interrogate type II FAS ACP-AT interactions. We utilize a covalent crosslinking probe to trap transient interactions between AcpP and FabD to elucidate the first x-ray crystal structure of a type II ACP-AT complex. Our structural data are supported using a combination of mutational, crosslinking, and kinetic analyses, and long timescale molecular dynamics (MD) simulations. Together, these complementary approaches reveal key catalytic features of FAS ACP-AT interactions. These mechanistic inferences suggest that AcpP adopts multiple, productive conformations at the AT binding interface, allowing the complex to sustain high transacylation rates. Furthermore, MD simulations support rigid body subdomain motions within the FabD structure that may play a key role in AT activity and substrate selectivity.

**Significance Statement:** The essential role of acyltransferases (ATs) in fatty acid synthase (FAS) and polyketide synthase (PKS) pathways, namely the selection and loading of starter and extender units onto acyl carrier proteins (ACPs), relies on catalytically productive ACP-AT interactions. Here, we describe and interrogate the first structure of a type II FAS malonyl-CoA:ACP-transacylase (MAT) in covalent complex with its cognate ACP. We combine structural, mutational, crosslinking and kinetic data with molecular dynamics simulations to describe a highly flexible and robust protein-protein interface, substrate-induced active site reorganization, and key subdomain motions that likely govern FAS function. These findings strengthen a mechanistic understanding of molecular recognitions between ACPs and partner enzymes and provide new insights for engineering AT-dependent biosynthetic pathways.

## INTRODUCTION

Fatty acid synthases (FASs) and polyketide synthases (PKSs) iteratively condense and often reduce ketide units to assemble compounds ranging from simple fatty acids to complex bioactive molecules.^1–4^ FASs and PKSs exist as either multidomain megasynthases (type I) or as discrete monofunctional enzymes (type II). They both share a common evolutionary origin,^3–7^ and therefore, FASs and PKSs often have related enzymatic components. Both systems require small, flexible acyl carrier proteins (ACPs) that are posttranslationally modified with phosphopantetheine (PPant) arms to shuttle covalently tethered reactive intermediates in the form of thioester bonds between catalytic partners. Catalysis is initiated by an acyltransferase (AT), which selects and loads starter and/or extender units onto the ACP. In type II FAS, extender units are loaded onto ACP via a malonyl-CoA:ACP transacylase (MAT), FabD, which catalyzes transfer of the malonyl moiety from malonyl-CoA to the PPant arm of *holo*-ACP to form malonyl-ACP (Figure 1). In contrast to type II FAS ATs, which only accept malonyl-CoA units, some ATs from PKSs accept a broader array of acyl-CoA units, as well as acyl-ACP units, increasing the structural diversity of their final products.^1,8–11^ Interestingly, various bacterial type II PKS gene clusters lack a canonical AT to form malonyl-ACP. This activity is instead provided by the endogenous FAS MAT.^12–14^ The central catalytic role of PKS and FAS MATs makes them attractive candidates for metabolic engineering and as targets for the development of new therapeutics.^15–20^

**Figure 1.**
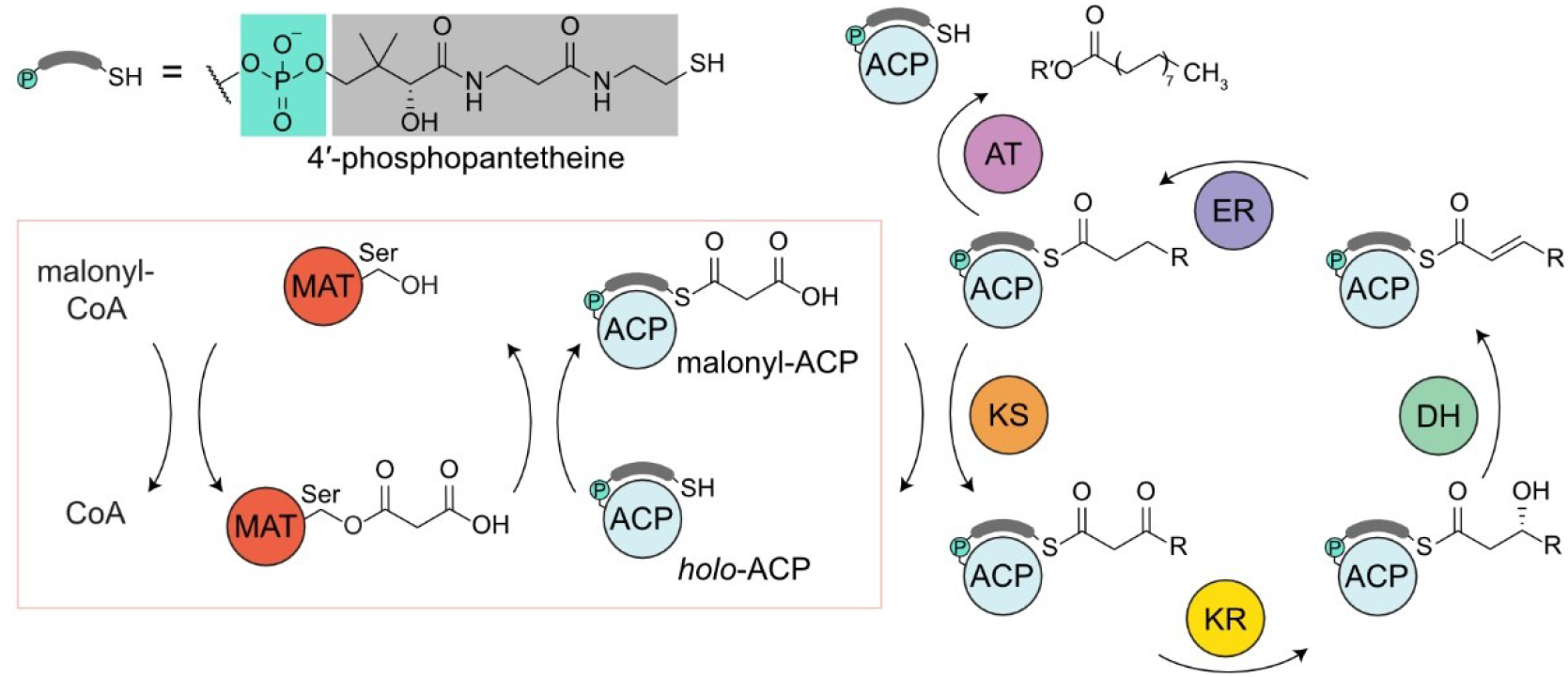
**Overview of saturated fatty acid biosynthesis from type II FAS,** highlighting the function of malonyl-CoA:ACP transacylases (MATs). The MAT (dark orange) is transiently acylated at an active site serine residue by malonyl-CoA. Then, the malonyl group is transferred to the 4′-phosphopantetheine arm of *holo*-ACP (pale blue). Additional downstream FAS enzymes catalyze chain extension, modification, and release from the synthase: the ketoacyl-ACP synthases (KS, orange), the NAD(P)H-dependent ketoacyl-ACP reductase (KR, yellow), the dehydratases (DH, aqua marine), the NAD(P)H-dependent enoylreductase (ER, purple) and the glycerol-3-phosphate acyltransferases (AT, lavender). The R and R’ groups represent the omitted portion of the acyl-ACP thioester and the glycerol 3-phosphate derived backbone of a phospholipid, respectively.

Engineering both type I and II FASs and PKSs is increasingly employed as a strategy to produce biofuels and bioactive natural products.^8,21–29^ In this context, ATs are of particular interest, as they act as metabolic gatekeepers that can direct the selection of key, biosynthetic building blocks. However, engineering efforts often lead to attenuated enzymatic activities^26,30^ or poorly soluble chimeras,^31^ likely due in part to an incomplete understanding of the transient nature of specific protein-protein interactions (PPIs) central to the iterative nature of FAS and PKS biosynthetic pathways. Interestingly, the high catalytic efficiency of FabD from *Escherichia coli*, which turns over substrate at nearly the diffusion limit (Figure S1, *k*_cat_/*K*_M_ ~10^8^ s^−1^ M^−1^),^32,33^ makes this MAT a particularly attractive target to create a diverse collection of acyl-ACPs for type II FAS and PKS engineering.^34,35^

Descriptions of type II FAS and PKS ACP-MAT interfaces benefited by docking simulations, NMR techniques, and binding affinity assays.^14,36,37^ While these studies suggest that MATs use shared binding motifs for malonyl-CoA and *holo*-ACP recognition, they often support different ACP-MAT binding interfaces. Recently, the x-ray crystal structures of two ACP-*trans*-AT covalent complexes from the vicenistatin and disorazole type I PKS pathways were elucidated.^38,39^ The ACP-AT binding modes and respective interface interactions in these structures describe notable differences, and it remains unclear whether the molecular underpinnings of PKS ACP-*trans*-AT interactions are fully shared with type II ACP-MAT interactions.

To address this, we utilized active site-selective crosslinking to covalently capture *E. coli* FabD in association with its cognate ACP, AcpP, to facilitate the elucidation of the first x-ray crystal structure of a type II FAS ACP-AT complex (PDB ID: 6U0J, 1.9 Å). We conducted biochemical assays that indicate the AcpP-FabD interface tolerates mutations to residues that initially appeared to support a specific and energetically stable protein-protein complex. In addition, we computationally probed this complex using long timescale molecular dynamics (MD) simulations. These computational studies support unanticipated dynamism and plasticity at the AcpP-FabD interface, likely explaining FabD’s tolerance to interface mutations with modest effects on turnover rates. A total of 32 μs of MD simulation data provides the most comprehensive sampling of AT dynamism to date, revealing rigid body subdomains motion, substrate-induced active site reorganization, and plasticity at the interface between the carrier protein and AT that likely explain the high catalytic rates of *E. coli* FabD. Interestingly, these domain motions have been proposed to play a role in the substrate promiscuity of the murine type I FAS AT.^40,41^ Our findings offer insights into the structures and dynamics of ACP-AT recognition in type II FASs. Collectively, these new results open up additional possibilities and challenges directed towards the design of protein-protein interfaces that support AT-mediated catalysis for the engineering of novel metabolic outputs.

## RESULTS AND DISCUSSION

### Crystallized crosslinked complex

In order to characterize the PPIs between a type II FAS ACP and its cognate AT, we crystallized a chemoenzymatically trapped complex of *E. coli* AcpP and its cognate AT, FabD. Classically, trapping covalent complexes between an ACP and a partner enzyme (PE) has been achieved through the synthesis of pantetheinamide crosslinking probes that react selectively and uniformly with the active site residues of the PE.^42–44^ These probes can be loaded onto an ACP via a one-pot chemoenzymatic method^45^ to produce *crypto*-ACPs, which, when mixed with cognate PEs, form crosslinked complexes.^46–49^ Despite recent successes developing active site fluorescent probes for ATs,^50^ targeting the active site serine of FabD with complementary pantetheinamide probes has proved challenging.

To produce sufficient quantities of crosslinked complexes for x-ray crystallography, we utilized a FabD S92C mutant that reacts with thiol-reactive pantetheinamide probes.^39,50,51^ Specifically, we generated C2-α-bromo-*crypto*-AcpP from *apo*-AcpP (Figure S2),^45^ which was subsequently incubated with FabD S92C to produce a crosslinked AcpP-FabD complex (where the “-“ denotes a covalent crosslink between the two proteins). This AcpP-FabD complex yielded diffraction quality crystals, resulting in the elucidation of a refined 1.9 Å-resolution structure (Table S1). The complex crystallized in the C2 space group and the asymmetric unit contains one molecule of FabD crosslinked to a single AcpP.

### Analysis of the 1.9 Å AcpP-FabD structure

The refined structure displays exceptional electron density for FabD, AcpP, and the synthetic crosslinker. FabD comprises two subdomains, a larger α/β hydrolase (ABH) subdomain spanning residues 1 to 125 and 203 to 308, and a smaller ferredoxin-like (FL) subdomain spanning residues 126 to 202 (Figure 2a). The active site of FabD is accessed along a large groove at the interface of the two subdomains which also serves as the PPant/CoA binding site. AcpP is bound between the two FabD subdomains abutting the aforementioned groove. In total, the AcpP-FabD interface buries 350 Å^2^ of surface area, making it the smallest reported AcpP-PE interface to date.^44^ The PPant arm extends from Ser36 of AcpP into the active site of FabD, where the α-carbon of the acetamide portion of the probe covalently attaches to the Cys92 mutant residue. The PPant arm makes a significant number of hydrophobic interactions derived almost exclusively from the FL subdomain of FabD, with the exception of Val280 from the ABH subdomain. Gln166, Asn160, and Asn162 line the floor of the groove, forming hydrogen bonding interactions with the PPant hydroxyl and amide functional groups (Figure 2d). A sulfate ion from the crystallization buffer is bound in the active site and coordinates to the conserved Arg117 residue that forms a bidentate interaction with the terminal carboxylate of the malonyl substrate (Figure S3a).^52^ The presence of the sulfate ion, the non-native thioether crosslinked bond, and the C-S bond length cause the carbonyl group of the probe to rotate away from the oxyanion hole formed by the backbone amides of Gln11 and Leu93 (Figures S1, S3b), which instead accepts two hydrogen bonds from the N_δ_ of Asn160 and the N_ε_ of His202.

**Figure 2.**
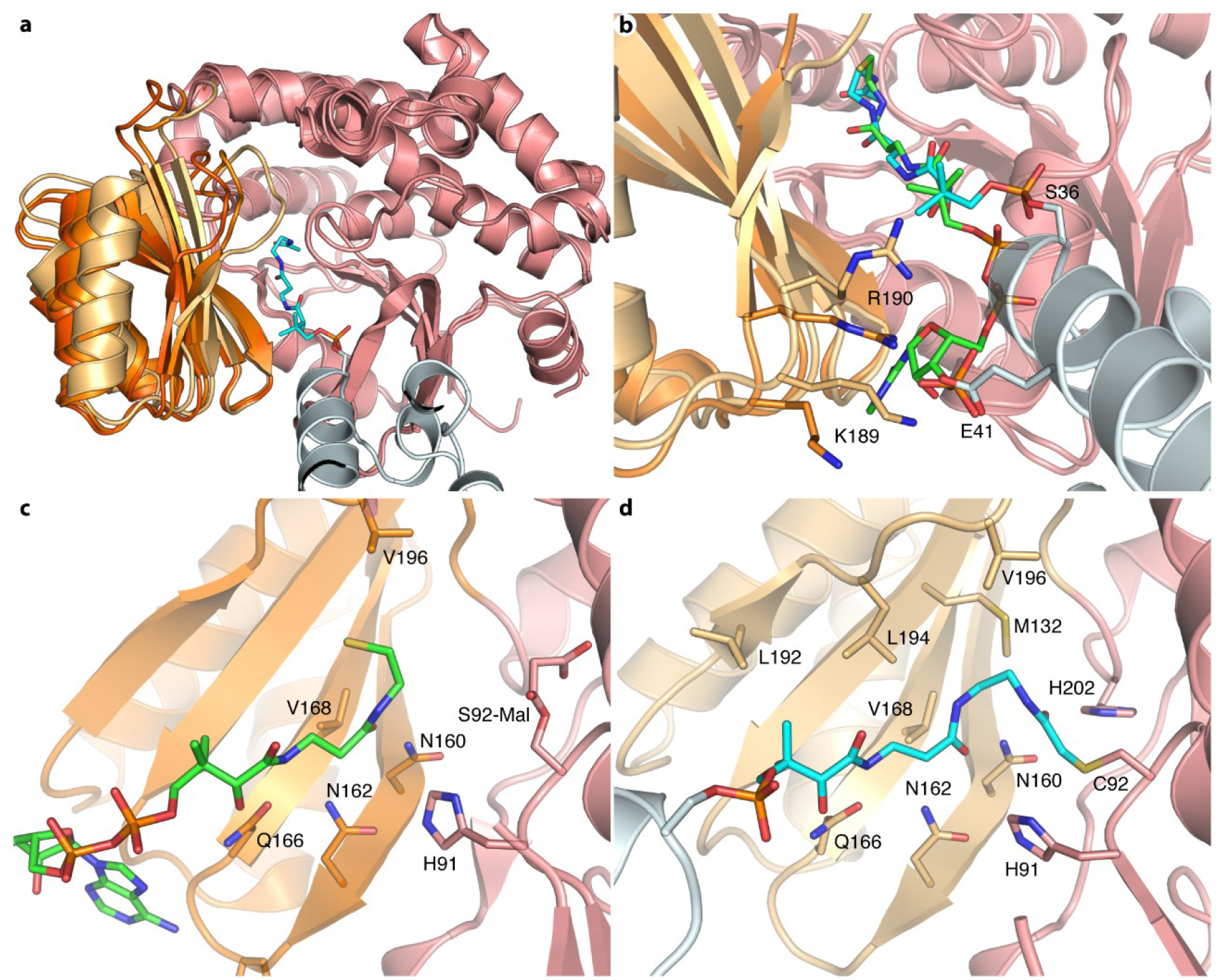
Ferredoxin-like subdomain movements and PPant interactions. **a**) Superposition of *apo*-FabD (dark orange, PDB ID: 1MLA), malonyl-CoA•FabD (orange, PDB ID: 2G2Z), and AcpP-FabD (light orange, PDB ID: 6U0J) over only the ABH subdomain (salmon) demonstrating the rigid body motion of FL subdomain with respect to ABH subdomain. **b**) Overlay of malonyl-CoA•FabD (orange) and AcpP-FabD (light orange) highlighting the similarities in interactions between CoA and *holo*-AcpP. The PPant arm (carbons colored cyan) and CoA (carbons colored green) are found in similar positions and conformations, and the 3′-phosphate of the CoA ribose ring is found in the same position as Glu41 of AcpP. **c**) Interactions between CoA and FabD in malonyl-CoA•FabD complex (PDB ID: 2G2Z). **d**) Interactions between the PPant arm and FabD in AcpP-FabD structure. A 4 Å distance cut-off was used to display FabD interacting residues in panels **c** and **d**. Only the interactions with the FL subdomain are shown for clarity in panel **d.**

The FL and ABH subdomains have been proposed to act as separate rigid bodies, with the FL subdomain capable of undergoing *en bloc* motions with respect to the larger ABH subdomain.^40,53^ Studies suggest FL subdomain mobility may be in part responsible for the reported substrate promiscuity of the AT domain from murine FAS.^40,41^ Given that FabD has undergone extensive structural characterization, we looked for potential changes in subdomain conformations induced by AcpP binding by performing a superposition using just the large ABH subdomains of *apo* FabD (PDB ID: 1MLA), malonyl-CoA-bound FabD (PDB ID: 2G2Z), and our AcpP-FabD crosslinked structure (PDB ID: 6U0J) (Figure 2a). Indeed, these structural overlays show significant variations in the relative positioning of the two subdomains. The FL subdomain possesses a 5 Å, hinge-like displacement toward the ABH subdomain when comparing the *apo*-FabD and AcpP-FabD structures (Figures 2a, S4). The most notable structural changes induced by these subdomain motions are found in the loop exiting the β4 strand of the FL subdomain, referred to herein as the β4-loop. This loop translates to cover the top of the FabD active site and PPant binding tunnel, and forms additional interactions between the two subdomains and the PPant moiety that are not present in either *apo*-FabD or malonyl-CoA-FabD structures (Figure S5). Interestingly, the dihedral angles of residues in the β4-loop only differ at Ser197 and Val198 when comparing *apo*-FabD, malonyl-CoA-bound FabD, and AcpP-FabD, suggesting that movement of this loop is largely dictated by the FL subdomain rigid body motion.

The FL subdomain, including the β4-loop, in the malonyl-CoA-bound FabD structure exists in an intermediate conformation between the *apo*-FabD and AcpP-FabD structures. This is likely due to the positioning of the free thiol from the hydrolyzed CoA moiety present in the active site that would sterically clash with Leu194 and Val196 on the β4-loop. Despite these differences in conformation, the PPant arm of CoA and AcpP overlay almost identically in the two structures and form a conserved hydrogen-bonding network with Gln166, Asn160 and Asn162 along the floor of the PPant binding tunnel (Figure 2b-d). It is worth noting that mutation of conserved residues along the β4-loop that precede the catalytic histidine (His 201 in FabD) results in changes in substrate specificities for type I PKS ATs. This may indicate that domain motions and the repositioning of the β4-loop over the active site play key roles in determining AT substrate selectivity and activity.^11,30,54,55^

### Analysis of the AcpP-FabD Interface

Here we describe the first type II FAS ACP-AT complex solved to date. Similar to all crosslinked AcpP-partner protein structures available,^46–49^ electrostatic complementarity between AcpP and FabD facilitates molecular recognition, (Figure S6) with the negatively charged AcpP interacting with a positive patch on FabD (Figure 3b) that likely drives initial association of AcpP and FabD. However, the AcpP-FabD crosslinked structure also departs from canonical FAS ACP binding motifs previously reported.^44^ It reveals a unique set of interfacial interactions between the two proteins involving the small helix on loop 1 of AcpP (α1’) and the N-terminal portion of helix II (α2) (Figure 3c-e, S7). Additionally, unlike the dehydratases (FabA and FabZ) and elongating ketosynthases (FabB and FabF) in *E. coli* FAS, FabD makes no discernable contacts with helix III (α3) of AcpP (Figure S6). Notably, α3 interactions likely facilitate chain flipping^56^ of AcpP-tethered cargo from the hydrophobic core of AcpP to the active sites of FabA, FabB, FabF and FabZ.^46,48,57^ Since the PPant arm of FAS *holo*-ACPs does not sequester within the helical hydrophobic core (PDB IDs: 5H9H, 3GZM, 2M5R, 2FQ0), the chain flipping process required for catalysis in these latter acyl-ACP•PE complexes is likely not operative in the delivery of the PPant arm of *holo*-ACP to the FabD active site.^58–61^ This may explain the lack of interfacial interactions involving α3 in the crosslinked AcpP-FabD structure.

**Figure 3.**
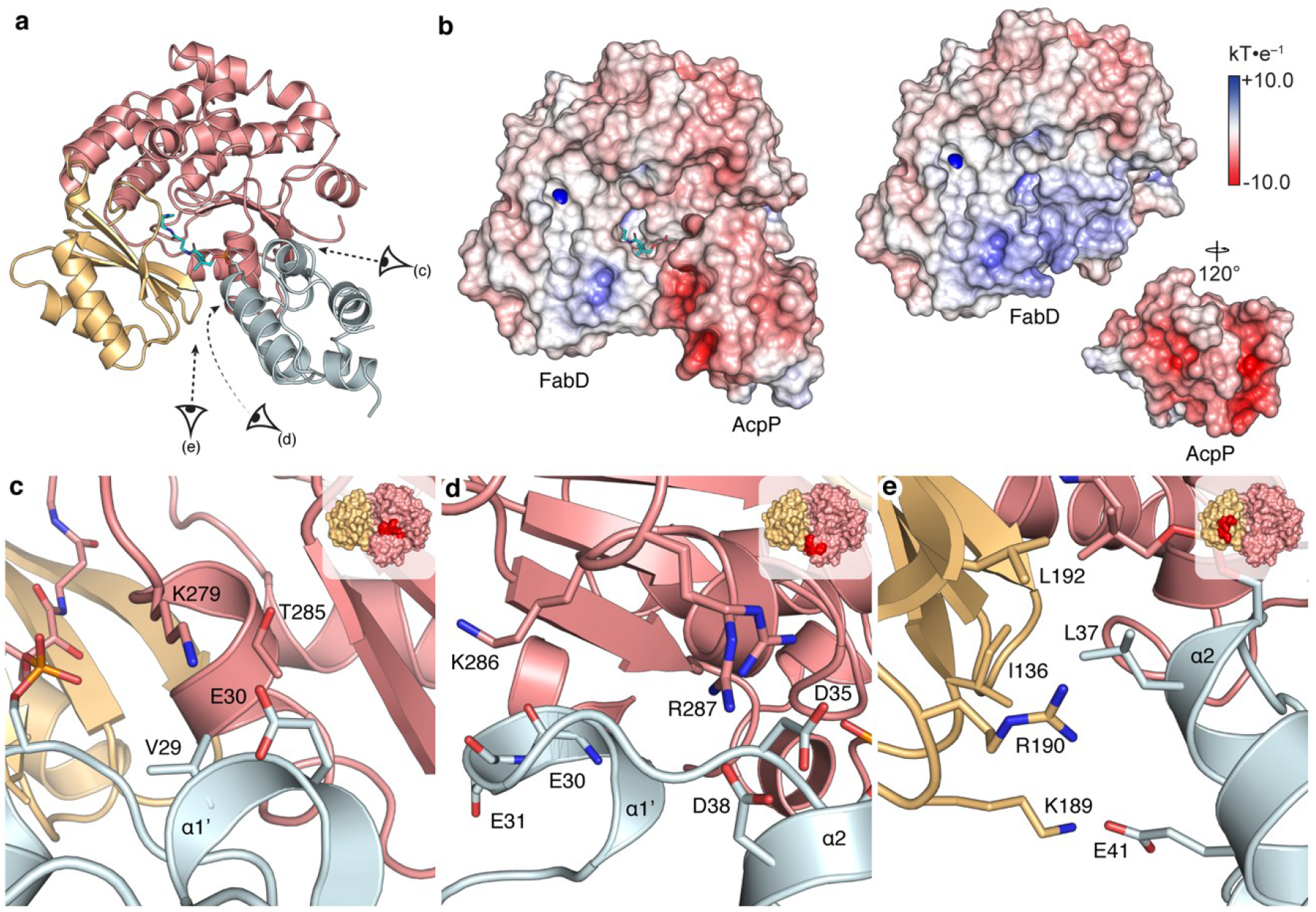
AcpP-FabD interface. **a**) Overview of the AcpP-FabD complex showing perspective eyes for panels c-e. The ABH and FL subdomains of FabD, as well as AcpP, are shown in salmon, light orange and light blue, respectively. **b**) AcpP-FabD, FabD, and AcpP electrostatic potentials mapped onto protein surface. **c**) Region 1 of FabD-AcpP interface. **d**) Region 2 of FabD-AcpP interface. Arg287 was modelled in two conformations, and both are shown. **e**) Region 3 of FabD-AcpP interface. Arg190 delineates into two conformations, but only the conformation closest to AcpP is shown for clarity. A 4 Å distance cut-off was used to display AcpP-FabD interacting residues in panels c-e, and the location of specific regions on FabD are shown as insets in the top-right corner of each respective panel.

In accordance with literature precedent,^39,49,51,62–65^ we investigated PPIs for AcpP-FabD by comparing the crosslinking efficiencies of a panel of FabD S92C interface mutants with AcpP loaded with a C2-α-bromo-pantetheinamide probe (Figure S8,S9b-c). Given that the FabD component of the interface is comprised almost exclusively of basic residues (Figure 3c-e), we generated three classes of FabD interface mutants: charge-neutralizing mutants (Arg to Ala or Lys to Ala), charge-conserving mutants (Arg to Lys), and charge-swapping mutants (Arg to Glu or Lys to Glu).

The location of specific interactions of the AcpP-FabD interface can be delineated into three regions (Figure 3c-e). Region 1 comprises the C-terminal portion of the ABH subdomain, which forms interactions with α1’ of AcpP. Lys279 of FabD forms a salt bridge with Glu30 of AcpP, and Val29 of AcpP fits into a hydrophobic pocket created by Lys279, Thr282, and Gly283 of FabD (Figure 3c). Both the charge-neutralized S92C/K279A and charge-swapped S92C/K279E mutants were generated to test the importance of this residue (Figure S9c). Both mutants show a considerable decrease in crosslinking efficiency, with 29% and 9% relative to S92C, for S92C/K279A and S92C/K279E, respectively.

Region 2 consists of Lys286 and Arg287, two conserved residues among bacterial type II MATs (Figure S9a). The key interactions in this region are between Arg287, modelled in two conformations, with both Asp35 and Asp38 at the top of AcpP’s helix II flanking the PPant Ser36 (Figure 3d). Interestingly, the guanidinium group of Arg287 forms cation-π stacking interactions with the adenine ring in the malonyl-CoA-bound FabD structure (PDB ID: 2G2Z), demonstrating its importance for both AcpP and malonyl-CoA binding.^14,36,37,52^ We produced all three classes of mutations for Arg287, resulting in charge-neutralized S92C/R287A, charge-swapped S92C/R287E, and charge-conserved S92C/R287K FabD mutants. Since Lys286 does not form a salt-bridge with AcpP in our structure, only the charge-neutralized mutant (S92C/K286A) was generated in this study. This mutant demonstrated a decrease in crosslinking efficiency, 23% relative to FabD S92C. The crosslinking efficiency of S92C/R287A was 14% relative to FabD S92C, while the charge-swapped mutant crosslinking efficiency decreased to 7%. Interestingly, crosslinking efficiency was partially restored to 36% of FabD S92C levels in the charge-conserved R287K mutant. (Figure S9c).

Region 3 consists of interactions between the FL subdomain and AcpP. In this region, Leu37 of AcpP sits in a hydrophobic groove formed by Ile136 and Leu192, and Glu41 from AcpP’s α2 forms a salt bridge with Lys189 of FabD (Figure 3e). Interestingly, overlays of AcpP-FabD and malonyl-CoA-bound FabD (PDB ID: 2G2Z) place the carboxylate side chain of Glu41 in the same location as the 3′-phosphate of the CoA ribose, indicating that CoA and AcpP form related interactions with FabD’s interface region 3 (Figure 2b). Despite the similarity in position, the CoA 3′-phosphate coordinates with Arg190 instead of Lys189, which is rotated away in the CoA-bound structure. Sequence alignments reveal that Lys189 is conserved among bacterial type II MATs, whereas Arg190 is conserved among both bacterial type II and eukaryotic mitochondrial MATs (Figure S9a). It is worth noting that the electron density for the side chains of residues Glu41, Lys189, and Arg190 is not well-defined. Glu41 and Lys189 have high B-factors, and Arg190 resides in two conformations, which suggests that the side chains of these residues are dynamic. In total, these data indicate that both Lys189 and Arg190 form interactions with Glu41 of AcpP. To evaluate the contribution of the FL subdomain to the AcpP-FabD interface, we mutated Leu192, Lys189 and Arg190 (Figure S9c). The S92C/L192A, S92C/K189A, and S92C/R190A mutants crosslinked with relative efficiencies of 13%, 62%, and 35%, respectively, compared to S92C. The charge-swapped mutant S92C/K189E showed minimal crosslinking (2%). In summary, results from these crosslinking studies validate the structural conclusions and indicate that the FL subdomain of FabD is critical to AcpP binding.

### Structural comparison with *trans*-AT-ACP interfaces

To date, only two other AT-ACP covalent complexes have been reported and analyzed, namely the *trans*-AT VinK in a covalent complex with VinL,^38^ and a second *trans*-AT from the disorazole synthase (DSZS) in a covalent complex with DSZS ACP1.^39^ *Trans*-AT PKSs do not contain a *cis*-acting AT domain within their modules to deliver the extender units onto the ACP, but instead, standalone ATs serve this role. VinK transfers a dipeptide intermediate between two ACPs (VinL and VinP1LdACP on the loading domain of module 1) during vicenistatin biosynthesis. Disorazole synthase is initiated by its *trans*-acting AT which malonylates the ACP of module 1. Elucidation of the AcpP-FabD complex allows analyses of the common features and differences of ACP recognition between modular type I PKS *trans*-ATs and type II FAS ATs. The overall subdomain organization of the three ATs, FabD, VinK, and DSZS AT (26.3% and 39.4% sequence identity with FabD, respectively) is similar, but the orientation of the ACPs notably differ between the three complexes. (Figure 4a-c). As observed in the AcpP•FabD complex, electrostatic contacts are important for both ACP•*trans*-AT complexes; however, hydrophobic interactions are more prominent features at the DSZS ACP1•DSZS AT and VinL•VinK interfaces than at the AcpP•FabD interface (Figures 4d-f, S10-11).^38,39^ The differences observed between these structures calls for a deeper look into the structural elements of the AT enzyme class.

**Figure 4.**
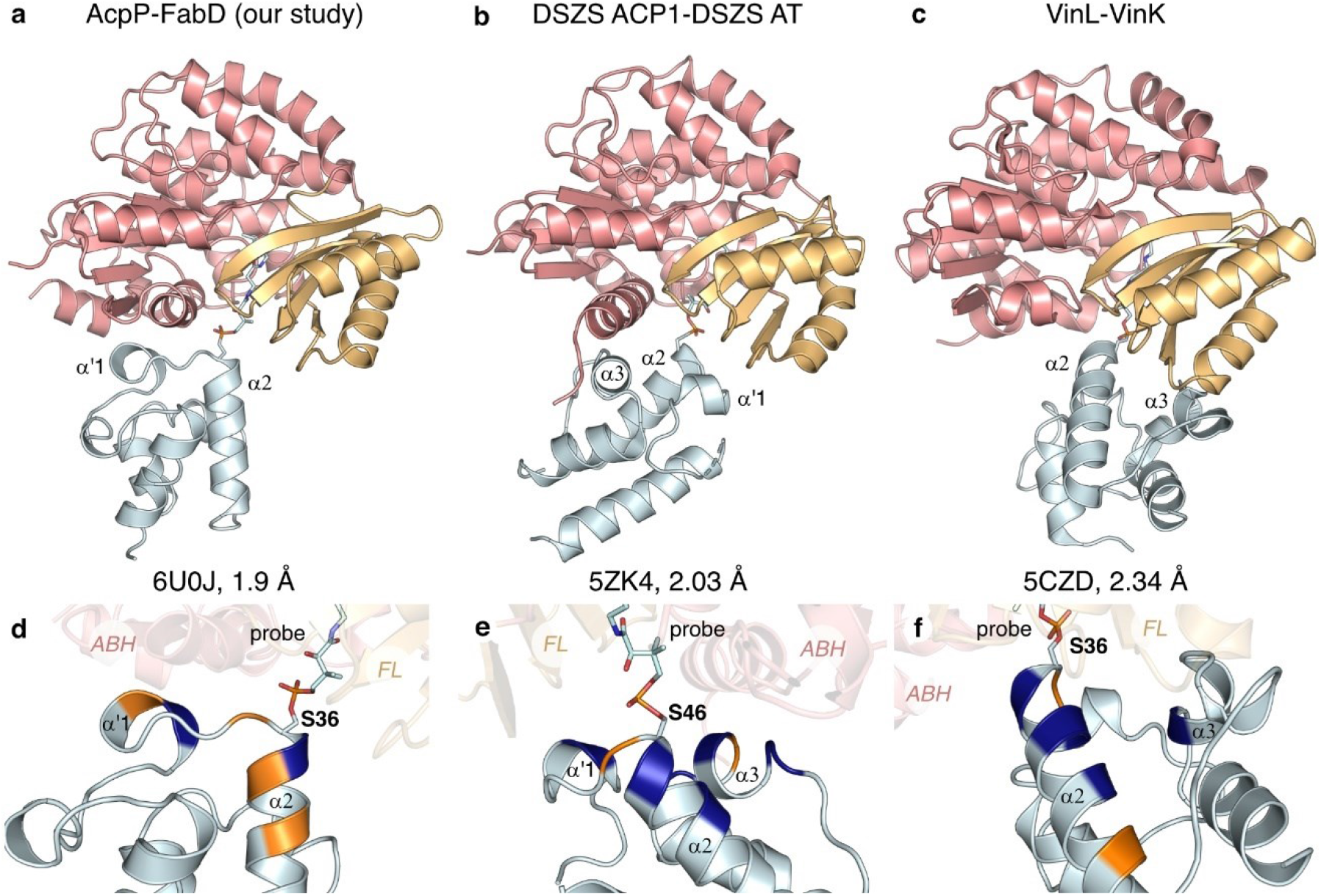
Comparison of the AcpP-FabD complex with the ACP-*trans*-AT complexes DSZS ACP1-DSZS AT and VinL-VinK. **a-c**) Cartoon representation of the AcpP-FabD, DSZS ACP1-DSZS AT and VinL-VinK complexes. The three complexes were aligned on FabD to better visualize the differences in ACP orientation. The FL and ABH subdomains of each AT are shown in light orange and salmon, respectively. The ACPs are colored in light blue. **d-f**) Close-up views at the ACP-AT interface for the three complexes displaying the nature of the interactions: blue, hydrophobic; orange: electrophilic. The interacting helices, catalytic serine and crosslinking probes are shown for each ACP. Note that **e** was rotated 180 ° from the above view to better visualize the interactions.

### Effect of PPIs on FabD’s transacylation reaction

We next performed kinetic analyses of FabD mutants by using a continuous, coupled fluorometric assay *in vitro* (Figure S12).^66^ We assayed our panel of FabD interface mutants along with three additional AcpP mutants, E30A, D38A, and E41A (Table 1). These three AcpP residues participate in salt bridges across the interface of regions 1, 2, and 3, respectively (Figure 3c-e). All constructs were analyzed by SDS-PAGE and mass spectrometry (Figure S13, Table S2). To compare the effect of each interface mutant, the transacylation rates are expressed as a percentage of the transacylation rate of wt *holo*-AcpP by wt FabD. Two concentrations of *holo*-AcpP (wt or mutant) were used, ACP_50_ and ACP_200_ (50 and 200 μM), which represent the concentration of *holo*-AcpP at the *K*_M_ value and at saturation for the wt system (Figure S14).

**Table 1.** Quantification of the transacylation rates of AcpP-FabD interface mutants. Reaction conditions: 28 °C, 50 mM sodium phosphate pH 6.8, 1 mM EDTA, 0.5 mM TCEP, 2 mM α-ketoglutarate, 0.4 mM NAD^+^, 0.4 mM thiamine pyrophosphate, 15 mU/100 μL α-ketoglutarate dehydrogenase, and 0.02 nM FabD. [ACP]_50_ corresponds to 50 μM *holo*-AcpP and 50 μM malonyl-CoA; [ACP]_200_ corresponds to 200 μM *holo*-AcpP and 200 μM malonyl-CoA. The activity of each interface mutant measured at the fixed concentration of AcpP indicated and was normalized to the corresponding activity of wt FabD, with wt AcpP for each tested condition (*see S.I. Relative activities*). Regions 1, 2 and 3 are shown in crimson, red, and orange, respectively. Experiments were run at least in triplicate, and data shown are reported as mean ± standard deviation.

| FabD | AcpP | Relative activity at [ACP]<sub>50</sub> (%) | Relative activity at [ACP]<sub>200</sub> (%) |
| --- | --- | --- | --- |
| wild type | wild type | 100.0 | 100.0 |
| K279A | wild type | 30.3 ± 6.9 | 23.9 ± 3.5 |
| K279E | wild type | 30.6 ± 7.6 | 30.2 ± 5.2 |
| wild type | E30A | 85.1 ± 7.9 | 99.5 ± 11.1 |
| K279A | E30A | 84.7 ± 10.3 | n.a |
| K279E | E30A | 14.2 ± 7.5 | n.a |
| R287K | wild type | 93.6 ± 3.2 | 102.7 ± 8.5 |
| R287A | wild type | 92.7 ± 9.4 | 89.0 ± 12.9 |
| R287E | wild type | 81.1 ± 7.8 | 99.6 ± 6.2 |
| wild type | D38A | 69.5 ± 4.8 | 83.2 ± 9.3 |
| R287K | D38A | 55.7 ± 5.4 | n.a |
| R287A | D38A | 77.3 ± 3.2 | n.a |
| R287E | D38A | 53.1 ± 6.9 | n.a |
| K286A | wild type | 63.1 ± 10.3 | 99.0 ± 10.5 |
| K286A/R287A | wild type | 96.8 ± 13.7 | 104.9 ± 13.9 |
| K286A/R287A | D38A | 77.9 ± 1.1 | n.a |
| K189A | wild type | 67.9 ± 7.3 | 77.4 ± 17.3 |
| K189E | wild type | 26.2 ± 4.8 | 104.0 ± 2 |
| wild type | E41A | 90.1 ± 1.9 | 95.8 ± 4.1 |
| K189A | E41A | 30.6 ± 2.1 | n.a |
| K189E | E41A | 10.6 ± 3.0 | n.a |
| R190A | wild type | 103.5 ± 9.5 | n.a |
| K189A/R190A | wild type | 40.1 ± 4.7 | 114.1 ± 8.7 |
| K189A/R190A | E41A | 15.2 ± 1.8 | n.a |
| L192A | wild type | 67.5 ± 8.2 | 101.0 ± 13.4 |

At ACP_200_, only three FabD variants show a decrease in activity compared to wt, namely K189A, K279A, and K279E. Overall, at ACP_50_, all tested FabD mutants have reduced activity compared to wt except for region 2 variants R287K and R287A that remain as active as wt, and R287E shows a slight loss in activity. We hypothesized that upon mutation of Arg287 to alanine, the fairly conserved Lys286 (Figure S9a) might have the rotational freedom to interact with Asp38 and compensate for the loss of the salt bridge between Arg287 and Asp38. We therefore tested the double mutant K286A/R287A, which remains as active as wt FabD for both AcpP concentrations. On the contrary, mutation of Lys279 in region 1 to either alanine or glutamate results in a larger decrease in relative activity as compared to FabD region 2 variants. Similarly, mutating Lys189 from region 3 also more significantly affects FabD activity, with a greater effect seen with the charge-swapped mutant, K189E as compared to the charge-neutralized variant K189A. To determine whether Arg190 participates in AcpP recognition despite its lack of discernable interactions in our AcpP-FabD structure, we evaluated the catalytic activity of R190A and K189A/R190A mutants. The R190A variant is similar in activity to wt, while the double mutant K189A/R190A is less active than K189A but faster than K189E. These data suggest that the interactions with the FL subdomain are mediated by both Lys189 and Arg190.

In addition to kinetically evaluating FabD interface residues, we also analyzed the transacylation rates of AcpP interface mutants, E30A, D38A, and E41A that coordinate interactions with FabD regions 1, 2, and 3, respectively. Interestingly, these mutations are less detrimental to FabD transacylation rates than their corresponding FabD interface mutants. In order to determine if the AcpP and FabD interface mutations were additive, we eliminated the three salt bridges at the interface by testing FabD interface mutants with their respective AcpP mutant counterparts. While assays of AcpP D38A and region 2 FabD mutants result in no more than a 50% drop in activity for all combinations tested, significantly slower transacylation rates were observed by combining mutant pairs from regions 1 and 3.

Results from all tested FabD and AcpP mutants demonstrate that the AcpP-FabD interface can tolerate mutations of residues that appear to be critical from the static crystal structure. We find that mutations on the FabD interface are more detrimental than their corresponding AcpP mutants, which may be due to the dynamic nature of AcpP. Our activity assays also show that mutating key residues on regions 1 and 3 of the FabD interface seem to influence the transacylation reaction more than region 2. Nevertheless, most mutations, even double mutants assayed with their corresponding AcpP mutant counterparts, do not abolish FabD’s activity.

### Dynamic analysis of the AcpP-FabD interface

The tolerance of the FabD interface to mutations, as indicated by our kinetic analyses, suggests that the interactions between AcpP and FabD are dynamic. Therefore, to further our understanding of the biomolecular processes that contribute to AcpP-FabD recognition, we subjected 17 protein assemblies, consisting of wt FabD and FabD mutants in the *apo*-, malonyl-CoA-, and malonyl-AcpP-bound states (Table S3), to MD simulations. We performed three independent 514 ns production-grade trajectories for each wt and mutant system, resulting in a total of 32.382 μs of simulation data. General analysis of these trajectories can be found in the SI: Molecular Dynamics Simulations Protocols, as well as in the supplementary Figures 15 to 22.

We analyzed trajectory data of the wt and mutant malonyl-AcpP•FabD complexes in order to propose a time-resolved understanding of the noncovalent interactions most critical for AcpP•FabD recognition. To do so, we identified all pairwise AcpP and FabD interactions that form over the course of the simulations using a 3.5 Å distance cutoff criterion (Figure 5). This analysis generally shows that the interactions at the AcpP•FabD interface are dynamic, and a number of interactions not observed in the x-ray structure are identified during the course of the simulations. Interestingly, none of the hydrophobic interactions identified in the AcpP-FabD structure were observed computationally (Figure 5). In contrast, the salt bridges identified in the x-ray structure are observed over the course of the simulations; however, they are in rapid exchange with an alternate set of salt bridges that are not resolved experimentally (Figure 5b). For example, the most frequently sampled salt bridge interaction between AcpP and FabD is a contact between Glu41 and Arg190 not present in the x-ray structure. Simulations of the malonyl-AcpP•FabD mutant complexes show the same general trend as the wt system and indicate that there are compensatory interactions that can be formed to stabilize these complexes despite the removal of key interface residues (Figure 5).

**Figure 5.**
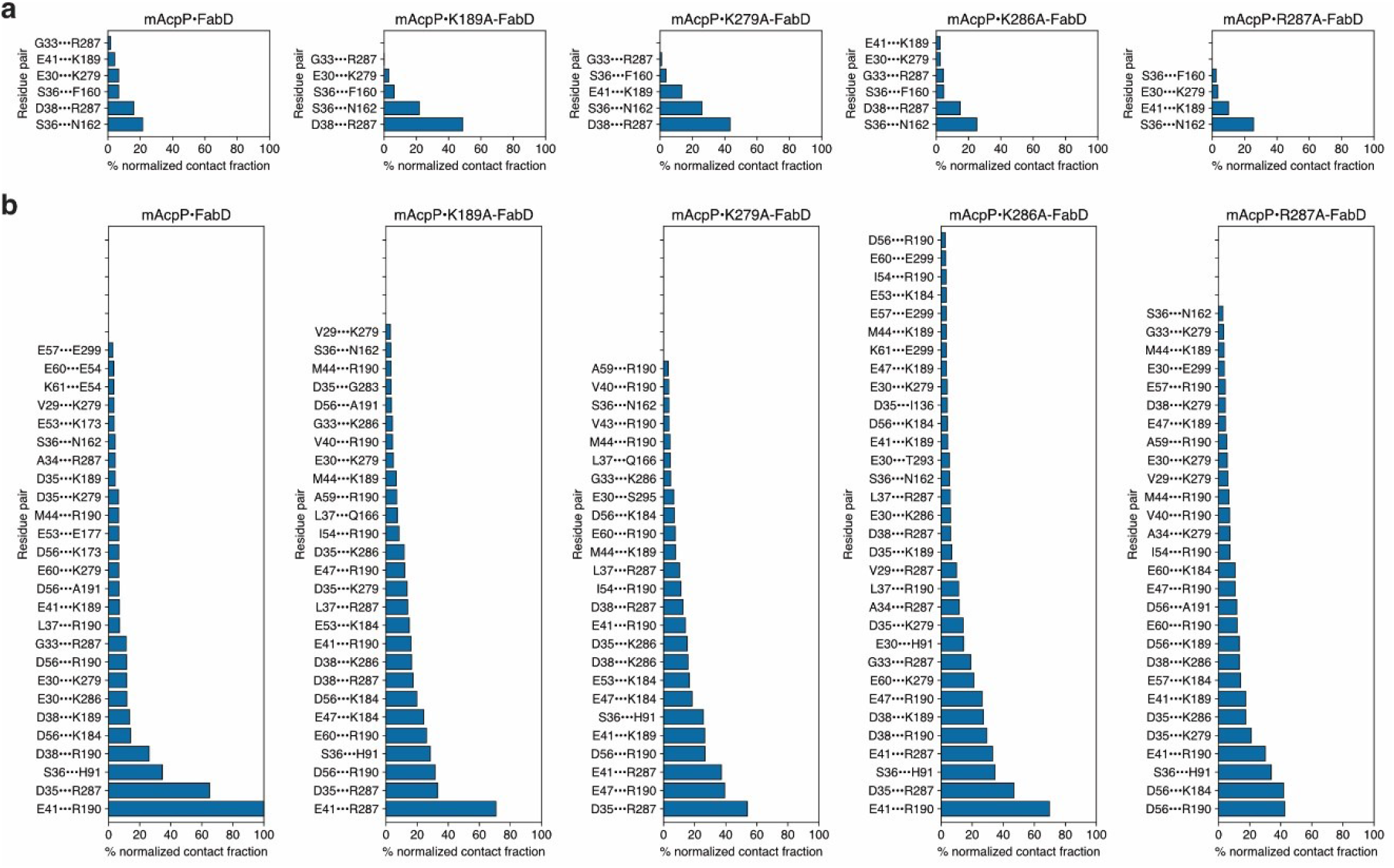
Time-resolved analysis of the contacts formed between FabD and its binding partner malonyl-AcpP. Contacts from the initial conditions and contacts formed during the course of the simulations are shown in panels **a** and **b**, respectively. Residues listed on the left side of the interaction pair correspond to AcpP and those on the right side of the pair correspond to FabD. The bar graphs indicate the normalized total contact fraction between FabD and malonyl-AcpP. The contact fraction is defined as the total fraction of simulation data in which a residue pair is engaged in an intermolecular contact. A distance criterion of 3.5 Å or less between a pair of heavy atoms defines such a contact. Only pairwise contacts with contact fractions (prior to normalization) greater than or equal to 0.10 are included in the plots above.

Given the rotational and translational motion of AcpP at the interface, we reasoned that these conformational changes may affect substrate organization in the FabD binding pocket. Interestingly, the contact between the malonyl carboxylate of malonyl-AcpP and the FabD active site Arg117, as well as the Ser92-His201 catalytic dyad distance, are maintained throughout the simulations despite the positional variation of AcpP (Figure S23-24). Taken together, these data suggest significant plasticity in the AcpP•FabD interface and provide a rationale for FabD’s tolerance to interface mutations, as AcpP can adopt multiple conformations at the FabD interface, while still properly positioning substrate in the active site for catalysis. In addition, our findings corroborate the smaller reduction of transacylation activity for AcpP mutations as compared to corresponding FabD mutations, as AcpP can rotate at the interface to form compensatory interactions (Figure 5).

### Substrate-induced reorganization of FabD active site

To determine the effect of substrate binding on active site organization, we performed a time-resolved analysis of interactions involving the conserved active site residues Ser92, Arg117, His201. The positively charged Arg117 residue has been shown to influence the preferences of ATs for α-carboxy-CoA substrates over α-descarboxy analogs. For example, the R117A mutant of FabD loads a number of α-descarboxy-CoA derivatives, including acetyl-CoA.^35^ Furthermore, Rangan and Smith showed that a similar Arg to Ala mutant of the malonyl-CoA/acetyl-CoA:ACP transferase of animal FAS results in a 100-fold decrease in its malonyl-CoA transferase activity, while its acetyl-CoA transferase activity is enhanced 6.6-fold.^67^ Studies by Dunn, Cane, and Khosla indicate that substrate discrimination occurs before the CoA substrate undergoes reaction at the AT active site and not by preferential hydrolysis of “misacylated” ATs after transacylation.^30^

MD simulations of *apo*-FabD suggest a possible mechanistic explanation for the broad role of Arg117 in regulating substrate selectivity in MATs. Analysis of simulation data of *apo*-FabD shows that in the absence of a substrate, the average distance between the Oγ of Ser92 and the center of geometry (COG) of Arg117’s terminal guanidinium moiety is 3.68 Å (Figure 6a). It must be noted that measuring from the arginine, histidine, or carboxylate COG reports distances larger than are typical of non-covalent interactions.^68^ Thus, simulation data shows that the dyad residues, Ser92 and His201, are not preorganized for catalysis in the *apo* state because Ser92 interacts stably with Arg117 (Figure 6a). Interestingly, simulations of *E. coli* malonyl-CoA•FabD show that α-carboxy-CoA substrate binding disrupts the Ser92-Arg117 interaction which in turn restores the Ser-His dyad (Figures S23-27). The average distance between the COG of Arg117 and the COG of malonyl carboxylate are 2.73 Å and 2.79 Å in simulations of wt malonyl-CoA•FabD (Figure S24) and wt malonyl-AcpP•FabD (Figure S27), respectively, suggesting a strong salt bridge interaction between substrate and enzyme in the bound state. With Arg117 engaged with the substrate, the catalytic dyad is restored, as the average distance between the Oγ of Ser92 and His201’s N_ε_2 sampled computationally is 3.40 Å and 2.95 Å for malonyl-AcpP•FabD and malonyl-CoA•FabD respectively, more than 1 Å shorter than what is observed over the course of simulations of *apo*-FabD (Figures 6a-c). These results suggest that substrate binding organizes the FabD active site for catalysis and provides an explanation for FabD’s (and MATs more broadly) preference for malonyl-CoAs or malonyl-AcpP over acetyl-CoA.^35^

**Figure 6.**
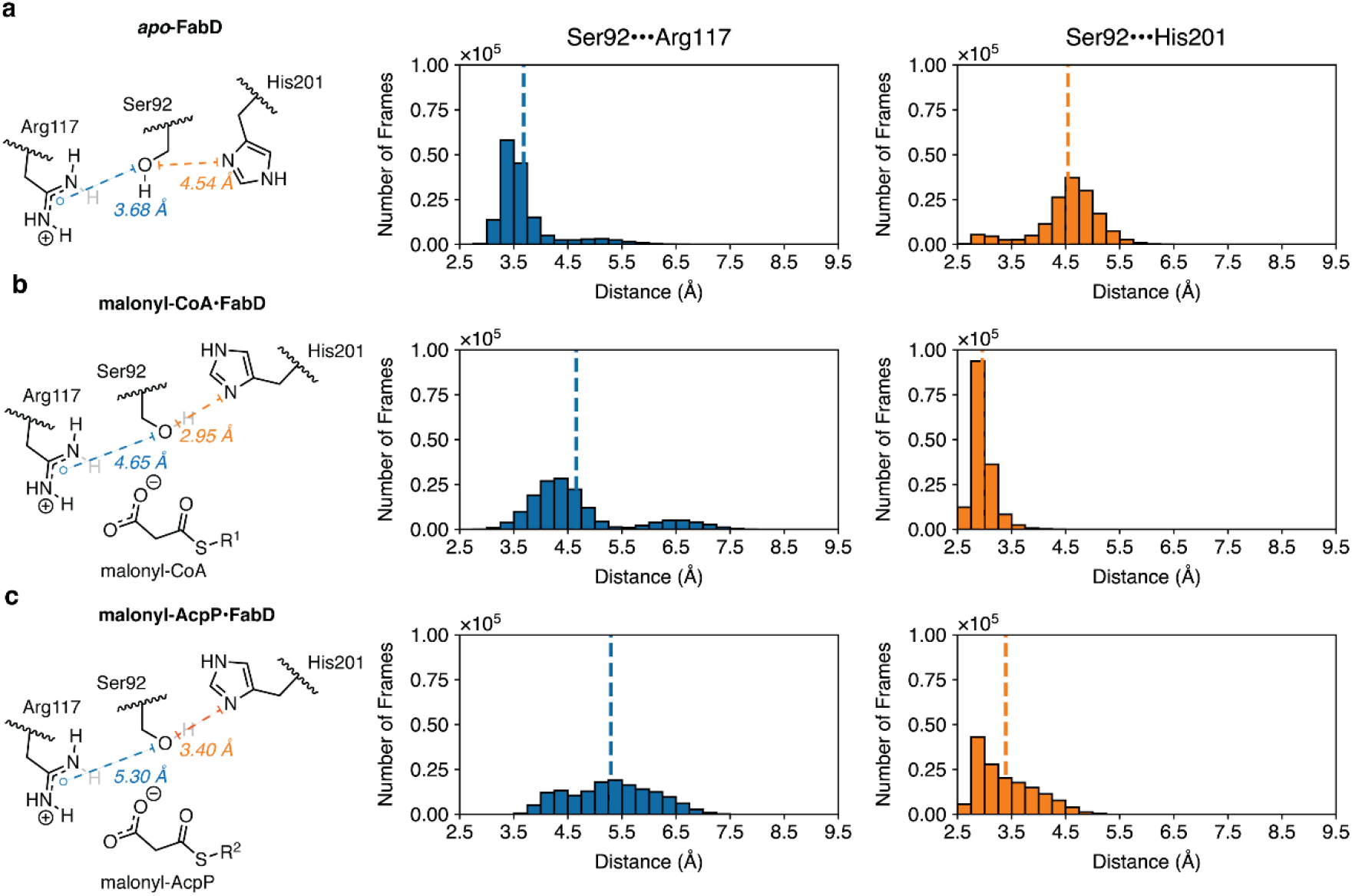
Analysis of key active site distances sampled in the *apo*- and substrate-bound states of FabD. Distribution of distances between two pairs of active site residues Ser92•••His201 and Ser92•••Arg117 sampled over the course of simulations of wt *apo*-FabD (**a**), malonyl-CoA•FabD (**b**). and malonyl-AcpP•FabD (**c**). The distance between Ser92 and His201 is measured using the Oγ of Ser92 and Nε of His 201, while the Ser92•••Arg117 distance is measured using the Oγ of Ser92 and the COG of Arg117’s guanidium moiety. Simulation data written every 0.5 ps were binned to prepare histograms using a bin width of 0.25 Å.

### Rigid body subdomain motions control the disposition of substrate-recognizing residues

Recent studies on the murine FAS AT have suggested that rigid body FL and ABH subdomain motion may play a role in determining AT substrate selectivity.^40,41^ Superpositions of our AcpP-FabD complex with *apo* and malonyl-CoA•FabD provide additional evidence for the occurrence of such motions (Figure 2a). Moreover, biochemical and structural data implicate residues found on the β4-loop of the FL subdomain, in particular those that immediately precede the catalytic histidine of ATs (His201 in FabD), play a role in determining AT preferences for α-branched *vs.* non-branched α-carboxy-CoA substrates.^11,30,54,55^ These residues (Val198, Ser199, and Phe200 in FabD) define part of the enzyme’s active site pocket and their steric bulk permits or prevents the binding of α-branched CoA substrates. We reason that rigid body motion of the FL subdomain, if present, may play a role in organizing the active site to preferentially accept either branched or non-branched CoA substrates. Therefore, we performed principal component analysis (PCA) to identify and highlight large-scale motions of the FL and ABH subdomains that may impact the activity and substrate specificity of FabD. Our analysis incorporates coordinate data from wt and mutants of *apo*-FabD (10.794 μs of simulation data), malonyl-CoA-FabD (9.252 μs of simulation data), and malonyl-AcpP (9.252 μs of simulation data).

Projections of the first principal component (determined via PCA) onto our coordinate data supports the hypothesis that the FL and ABH subdomains act as rigid bodies involving the displacement and rotation of the smaller FL subdomain away from ABH subdomain (Figure 7). That this motion is represented by the first principal component, the orthogonal component with the largest variance (*i.e.*, most dominant “mode” describing FabD motion), supports its role as an essential feature of FabD’s function. A comparison of the first principal components generated from the dataset described above indicates that, as suggested by the comparison of x-ray structures presented in Figure 2, the FL subdomain moves independently of the ABH subdomain (Figure 7, Movies S1-S3). The rigid body motions evaluated computationally disambiguate a motion of FL subdomain towards and away from the ABH domain to gate substrate access to and from the active site. In addition to active site gating, this rigid body motion also results in the movement of the β4-loop towards the base of the active site, not only organizing the His201 for catalysis, but also positioning the key substrate-sensing residues Val198, Ser199, and Phe200 to selectively interact with *E. coli* FabD’s preferred substrate (Movies S1-S3).

**Figure 7.**
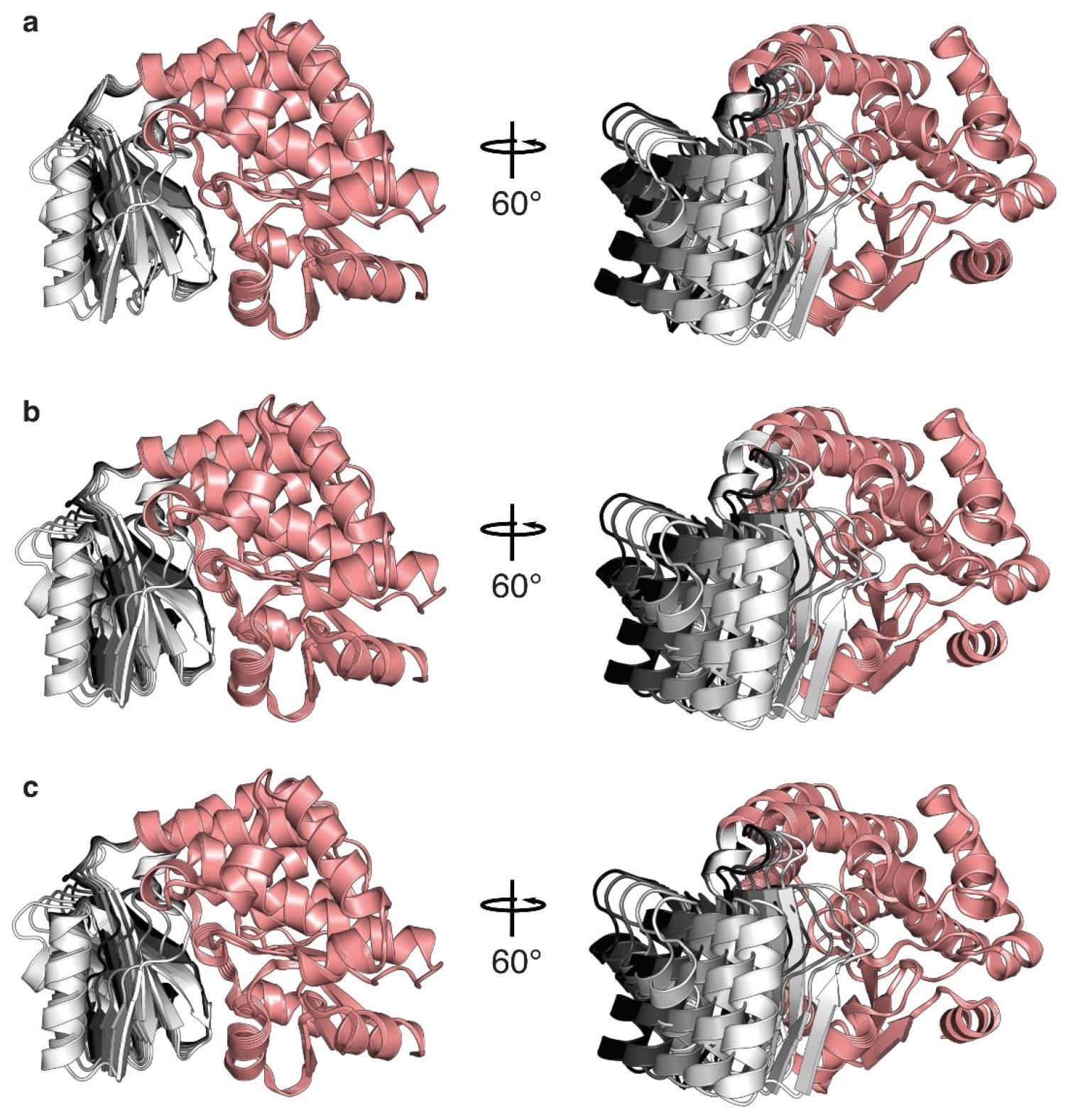
Projection of principal component analysis data into coordinate space. Overlaid snapshots of the pseudo-trajectories illustrating the 1^st^ mode of motion of the FabD monomer portions of **a**) *apo*-FabD, **b**) malonyl-CoA•FabD, and **c**) malonyl-AcpP•FabD. The ABH subdomains are colored salmon while the FL subdomains are colored based upon the distance between the FL subdomain, such that more darkly colored subdomains are further from the ABH subdomain.

## CONCLUSION

In this study, we use *E. coli* type II FAS as a structural and mechanistic model to better understand type II ACP-AT interactions and their catalytic role in fatty acid biosynthesis. Utilizing a pantetheinamide covalent crosslinking probe, we trapped and crystallized the *E. coli* AcpP in complex with FabD, allowing us to elucidate the first crystal structure of a type II ACP-AT complex at 1.9 Å resolution. The AcpP-FabD binding mode is different than previously described AcpP complexes and represents the smallest AcpP-PE interface reported to date. Despite the small number of experimentally determined interfacial interactions, mutation of these interface residues does not result in significant reductions in FabD transacylation activity. Long timescale MD simulations of AcpP in complex with wt FabD and FabD interface mutants demonstrate that AcpP can undergo considerable rotational and translational motion at the FabD interface while still properly positioning its PPant-tethered substrate for catalysis. These results are in line with kinetic analyses of FabD interface mutants and suggest a dynamic interface dominated by polar and charged interactions that are in rapid exchange. Furthermore, MD analysis of FabD in both its liganded and unliganded states corroborate the presence of *en bloc* rigid body motions of its FL and ABH subdomains and provide additional insights into Arg117’s role in substrate mediated reorganization of the FabD active site. Taken together, these findings represent a significant advancement in our understanding of type II AcpP-AT interactions, AT dynamics, and AT substrate selectivity. The structural, biochemical, and computational analyses of these systems reported herein represent another necessary step toward a broader and more complete understanding of PPIs in type IsI FAS and natural product biosynthesis.

## Materials and Methods

The details for all of the experimental protocols are described in SI Materials and Methods.

## Data deposition

The atomic coordinates and structure factors have been deposited in the Protein Data Bank, www.pdb.org (PDB ID code 6U0J).

## Acknowledgments

This work was supported by NIH GM095970 and GM31749. L.E.M. was supported by an Early Postdoc.Mobility Fellowship from the Swiss National Science Foundation. J.T.M. was supported by T32 GM832626. T.D.D. is a San Diego IRACDA Postdoctoral Fellow supported by NIH K12 GM0658524 and NIH K99 GM129454. Portions of the work were also funded by the Arthur and Julie Woodrow Chair at the Salk Institute (to J.P.N.) and the Howard Hughes Medical Institute (to J.P.N.). The authors thank Dr. Gordon Louie for assistance in x-ray data collection, processing, and refinement, Marianne Bowman for assistance with protein crystallization and the UCSD Chemistry and Biochemistry Mass Spectrometry Facility for the molecular weight determination of FabD and AcpP wt and variants. We further acknowledge the following organizations for support of the computer simulation work reported herein: NIH GM31749 (to J.A.M.) and the San Diego Supercomputer Center (to J.A.M.).

## Notes

### Competing Interest Statement

The authors have declared no competing interest.

