## Supplementary Information for "Structure and Dynamic Basis of Molecular Recognition Between Acyltransferase and Carrier Protein in *E. coli* Fatty Acid Synthesis"

**This PDF file includes:**

Supplementary text
Figures S1 to S28
Tables S1 to S4
Legends for Movies S1 to S3
SI References

**Other supplementary materials for this manuscript include the following:**

Movies S1 to S3

**Table of Contents**

**A. Biological Protocols**

**A1.** Expression and purification of FabD and AcpP wt and mutants
**A2.** FabD thrombin $His_6$-tag cleavage
**A3.** *Apo*-AcpP loading and C2-α-bromo-*crypto*-AcpP purification
**A4.** Production and purification of AcpP-FabD crosslink for crystal trials
**A5.** Crystallization, data collection, and refinement
**A6.** AcpP holofication for kinetic assay
**A7.** Site-directed mutagenesis
**A8.** α-ketoglutarate dehydrogenase coupled activity assay
**A9.** Analysis of kinetic data
**A10.** Michaelis-Menten kinetics
**A11.** Relative activities of FabD and AcpP mutants
**A12.** Preparation of C2-α-bromo-*crypto*-AcpP for analytical crosslinking efficiency assays
**A13.** Analytical crosslinking assays
**A14.** Time-dependent crosslinking

**B. Synthetic Protocols**

**B1.** General synthetic methods

**C. Molecular Dynamics Simulations Protocols**

**C1.** Structure preparation
**C2.** Simulation preparation
**C3.** Simulation methodology
**C4.** Analysis and visualization of simulation data

**D. Supplementary Figures and Tables**

**Figure S1.** Reaction catalyzed by FabD
**Figure S2.** Chemoenzymatic loading of *apo*-AcpP with C2-α-bromo-pantetheine and subsequent FabD S92C crosslinking
**Figure S3.** Comparison of AcpP-FabD and malonyl-CoA•FabD active site interactions
**Figure S4.** Measurement of FL subdomain rigid body motions
**Figure S5.** Interaction networks formed between FL subdomain and ABH subdomain in AcpP-FabD structure
**Figure S6.** AcpP Interactome
**Figure S7.** Overall organization of AcpP in the AcpP-FabD structure
**Figure S8.** Purified FabD S92C interface mutants for crosslinking validation
**Figure S9.** Validation of FabD•AcpP interface *via* mutagenesis and crosslinking
**Figure S10.** ACP-AT electrostatic potentials maps of ACP-AT complexes and respective monomers
**Figure S11.** Interfaces residues of trans-ATs compared to FabD
**Figure S12.** FabD continuous coupled assay using α-ketoglutarate dehydrogenase (KDH)
**Figure S13.** Coomassie-stained SDS-PAGE and urea-PAGE gels
**Figure S14.** Michaelis-Menten curves of wt FabD
**Figure S15.** $C_\alpha$ root mean square analysis (RMSD) in Ångstroms (Å) of the simulations of wildtype (wt) *apo*-FabD and five FabD mutants
**Figure S16.** $C_\alpha$ root mean square analysis (RMSD) in Ångstroms (Å) of the simulations of wildtype (wt) FabD and *apo*-FabD bound with malonyl-CoA performed using ff14SB/GAFF2 force field parameter

**Figure S17.** $C_\alpha$ root mean square analysis (RMSD) in Ångstroms (Å) of the simulations of malonyl-AcpP-bound wildtype (wt) FabD and *apo*-FabD
**Figure S18.** Heavy atom root mean square analysis (RMSD) in Ångstroms (Å) of the malonyl-CoA substrate of the wt and mutant malonyl-CoA•FabD complexes.
**Figure S19.** Heavy atom root mean square analysis (RMSD) in Ångstroms (Å) of the wt and mutant malonyl-FabD•FabD complexes
**Figure S20.** $C_\alpha$ root mean square analysis (RMSD) in Ångstroms (Å) of the simulations of wildtype (wt) *apo*-FabD* and *apo*-FabD**
**Figure S21.** Root mean square fluctuations (RMSFs) of each residue of FabD of the variants of the *apo*-FabD, malonyl-CoA•FabD, and malonyl-AcpP•FabD system subjected to MD simulation
**Figure S22.** Root mean square fluctuations (RMSFs) of each residue of FabD of the variants of the *apo*-FabD, malonyl-CoA•FabD, and malonyl-AcpP•FabD system subjected to MD simulation
**Figure S23.** Distribution of distances between key active site residues sampled throughout the course of MD simulations of malonyl-AcpP•FabD variants
**Figure S24.** Distribution of distances between key active site residues and malonyl portion of malonyl-AcpP sampled throughout the course of MD simulations of malonyl-CoA•FabD variants
**Figure S25.** Distribution of distances between key active site residues sampled throughout the course of MD simulations of *apo*-FabD variants
**Figure S26.** Distribution of distances between key active site residues sampled throughout the course of MD simulations of malonyl-CoA•FabD variants
**Figure S27.** Distribution of distances between key active site residues and malonyl portion of malonyl-AcpP sampled throughout the course of MD simulations of malonyl-CoA•FabD variants
**Figure S28.** Time-dependent C2α-bromo-*crypto*-AcpP-FabD S92C crosslinking

**Table S1.** X-ray crystallography data collection and refinement statistics.
**Table S2.** Molecular weight of FabD and holo-AcpP variants
**Table S3.** FabD systems subjected to molecular dynamics simulations
**Table S4.** PCR primers used in this study.

**E. $^{1}H$ and $^{13}C$ NMR Spectra**

**F. SI References**

### A. Biological Protocols

**A1. Expression and purification of FabD and AcpP wt and mutants.** The genes encoding for FabD and AcpP were inserted in a pET28b vector with a cleavable N-terminal $His_6$-tag. The proteins (wt and variants) were expressed in *E. coli* BL21(DE3) cells. For each variant, a single colony was selected on the agar plate and grown overnight at 37 °C in 5 ml Luria-Bertani (LB) medium supplemented with 50 µg/ml kanamycin (KAN). Then, 1 L of LB medium (50 µg/mL KAN) was inoculated with the 5 mL-LB preculture and incubated at 37 °C until an optical cell density of 0.7 was reached. After induction with 0.5 mM IPTG, cells were grown for 3 h at 37 °C. Cells were harvested by centrifugation (500 RCF, 30 min) and the pellet was stored at – 20 °C. Cells were resuspended in lysis buffer (50 mM Tris pH 8.0; 300 mM NaCl; 10 % glycerol) and lysed by sonication. After centrifugation (17.400 RCF, 45 min), the supernatant was transferred to Ni-NTA-column and washed with 10 CV of lysis buffer and 2 x 10 mL lysis buffer containing 10 mM imidazole. The proteins were eluted with elution buffer (lysis buffer containing 250 mM imidazole). All FabD and AcpP mutants expressed with yields comparable to the wt proteins, ranging from 25 ± 5 and 15 ± 5 mg of protein / L of LB, respectively. FabD wt and variants were dialyzed twice over 1 L of dialysis buffer at 4 °C (50 mM phosphate pH 6.8, 150 mM NaCl, 0.5 mM TCEP and 10 % glycerol), aliquoted, frozen in liquid nitrogen and stored at -80 °C. AcpP wt and variants (expressed as a mixture of *apo* and *holo* forms) were subjected to His-tag cleavage by thrombin and the resulting His-tag free AcpPs were purified using Ni-NTA-column supplemented with Benzamidine Separophore (bioWorld). The resulting *apo-/holo*-AcpPs were then converted to their *holo* form.

**A2. FabD S92C thrombin $His_6$-tag cleavage**. The $His_6$-tag of FabD S92C (purified as reported above) was cleaved using bovine thrombin (2 U per 1 mg protein) overnight at 6 °C while dialyzing against dialysis buffer (50 mM Tris, 150 mM NaCl, 10% glycerol, pH 8.0). Resulting solutions were re-purified using Ni-NTA resin (Thermo Fisher Scientific) to remove un-cleaved proteins. The resulting $His_6$-tag free FabD was further purified by FPLC using the HiLoad Superdex 200 (GE Biosciences) size exclusion column. The eluted protein was collected and concentrated to 2-4 mg•$mL^{-1}$ using Amicon Ultra Centrifuge Filters (MilliporeSigma) with 10 kDa molecular weight cut off.

**A3. *Apo*-AcpP loading and C2-α-bromo-*crypto*-AcpP purification.** C2-α-bromo-pantetheinamide crosslinkers were loaded onto *apo*-AcpP using a "one-pot" chemoenzymatic method.[1] This method utilizes the CoA biosynthetic enzymes (CoaA, CoaD, CoaE) to form CoA analogues and a phosphopantetheinyl transferase (PPTase, Sfp) to load them onto *apo*-AcpP, resulting in *crypto*-AcpP. Final reaction concentrations: 1 mg•mL$^{-1}$ *apo*-AcpP, 0.04 mg•mL$^{-1}$ Sfp, 0.01 mg•mL$^{-1}$ CoaA, 0.01 mg•mL$^{-1}$ CoaD, 0.01 mg•mL$^{-1}$ CoaE, 50 mM potassium phosphate pH 7.2, 12.5 mM $MgCl_2$, 1 mM DTT, 0.2 mM C2-α-bromo-pantetheinamide, and 8 mM ATP. The stock solution of C2-α-bromo-pantetheinamide crosslinking probe was prepared in DMSO at a final concentration of 50 mM. Reactions were incubated at 37 °C for 16 h and then purified by HiLoad Superdex 75 (GE Biosciences) size exclusion column. The eluted protein was collected and concentrated using Amicon Ultra Centrifuge Filters (Millipore Sigma) with 3 kDa molecular weight cut off up to a concentration at 1-3 mg•mL$^{-1}$.

**A4. Production and purification of AcpP-FabD complex for crystal trials.** The crosslinking reaction was carried out by mixing C2-α-bromo-*crypto*-AcpP with $His_6$-tag free FabD S92C at 2:1 (C2-α-Bromo-*crypto*-AcpP:FabD S92C) ratio at 37 °C for 16 h. Reactions were analyzed by 12 % SDS PAGE. Crosslinked AcpP-FabD complex was purified by a Ni-NTA resin (Thermo Fisher Scientific), allowing the removal of uncrosslinked $His_6$-tag cleaved FabD S92C from the reaction mixture. The bound AcpP-FabD complex and was eluted from the column using 250mM imidazole. The resulting fractions containing the AcpP-FabD complex were purified on a HiLoad 16/600 Superdex 200 PG (GE Biosciences) size exclusion column using minimal buffer (12.5 mM Tris, 100 mM NaCl, pH 8.0). The resulting protein complex was greater than 95 % in purity, determined by SDS PAGE, and was concentrated to 8 mg•mL$^{-1}$ using Amicon Ultra Centrifuge Filters (MilliporeSigma) with 10 kDa molecular weight cut off. The concentrated crosslinked complex was immediately used for protein crystallization or flash-frozen and stored in –80 °C freezer for later use.

**A5. Crystallization, data collection, and refinement.** The crystals of the AcpP-FabD crosslinked complex were grown by vapor diffusion using the hanging drop method at 6 °C. In detail, 1 µL of crosslinked complex (8 mg•mL$^{-1}$) was mixed with 1 µL of corresponding mother liquor and the mixture was placed inverted over 500 µL of the well solution (hanging-drop method). The AcpP-FabD complex crystallized in 2.3-2.7 M $AmSO_4$, 1% PEG 400, and 0.1 M sodium cacodylate pH 6-7. These conditions initially yielded multiple yellow spherulites that eventually resulted in numerous small needle-like crystals in 1-2 weeks. The best well condition (2.5M $AmSO_4$, 1% PEG 400, pH 7.0) resulted in a few small ovular plates (diffracted to 1.9 Å). All data were collected at the Advanced Light Source (ALS) synchrotron at Berkeley. Data were indexed using iMosflm[2] then scaled and merged using the aimless program from the CCP4 software suite.[3] Scaled and merged reflection output data was used for molecular replacement and model building in PHENIX.[4] Phases were solved using molecular replacement using FabD (PDB ID: 1MLA) as a search model with program Phaser from the PHENIX software suite. AcpP (PDB ID: 2FAC) was manually placed and rebuilt in the residual electron density using Coot.[5] The parameter file for the covalently bonded 4'-phosphopantetheine was generated using eLBOW in the PHENIX software suite.[6] Manually programmed parameter restraints using values provided by Jligand (CCP4)[3,7] were used to create the associated covalent bonds between 4'-phosphopantetheine to Ser36 and Cys92 during refinement.

**A6. AcpP holofication for kinetic assay.** *holo*-AcpP (wt or mutants) was prepared as previously described.[8] Briefly, AcpP (*apo*/*holo*) was incubated at 37 °C overnight in the presence of the phosphopantetheinyl transferase Sfp from *Bacillus subtilis*. Final reaction concentrations were the following: 50 mM phosphate pH 6.8, 50 mM NaCl, 0.5 mM TCEP, 12.5 mM $MgCl_2$, 1 mM coenzyme A, 0.05 mg/mL Sfp and 2.2 mg/mL AcpP. The completion of the reaction was determined by urea PAGE (Figure S13, 2b) and mass spectrometry (Table S2). The resulting *holo*-AcpP was further purified on a Superdex 75 HiLoad 16/60 size exclusion chromatography (SEC) column equilibrated with buffer (50 mM sodium phosphate pH 6.8, 125 mM NaCl, 5 % (v/v) glycerol, 0.5 mM TCEP). The eluted AcpP was collected, concentrated using Amicon Ultra Centrifuge Filters (Millipore) with 3 kDa molecular weight cut off, frozen in liquid nitrogen and stored at -80 °C.

**A7. Site-directed mutagenesis**. FabD and AcpP mutants (except FabD S92C and AcpP D38A) were generated by polymerase chain reaction (PCR) amplification process using the primers listed in Table S4. FabD S92C and AcpP D38A constructs were provided from previous studies.[9,10] While D35 interacts with R287 on our structure (Figure 3), it is part of the highly conserved DSL motif at the N-terminus of helix II that serves as a recognition sequence and the site of PPant modification by ACPS, the *holo*-AcpP synthase in *E. coli*.[11] Mutating D35 would result in the *apo* form of any AcpP variant and was therefore not further investigated.

**A8. α-Ketoglutarate dehydrogenase coupled activity assay.** The KDH coupled assay (Figure S12) used in this study was adapted from previous work.[12–14] Briefly, all enzymatic reactions were in a final volume of 100 µl and performed in 96-well half-area plates (Corning, Fisher Scientific).

NADH fluorescence was monitored using a Varioskan Lux microplate reader (Thermo Scientific) at the excitation and emission wavelengths of 340 and 460 nm, respectively. The background reaction rate of this assay was determined in the absence of FabD and subtracted to each of the reactions performed in the presence of FabD. Equidistant kinetic measurements were taken every 5 s for 5 min at 28 °C.

**A9. Analysis of kinetic data.** Kinetic data were analyzed with Origin software. Data points from 20–120 s were fitted by linear regression to give the initial velocity of substrate turnover. The averages of triplicates were calculated, and the background reaction rate subtracted. Relative fluorescence units (RFU) were converted into concentrations (µM) using a calibration curve (Figure S12).

**A10. Michaelis-Menten kinetics.** The Michaelis-Menten parameters of FabD transacylation reaction have been previously reported.[12,15,16] However, to ensure that we would perform our mutagenesis analysis at the expected AcpP concentrations (1x $K_M$ (wt AcpP) and 4x $K_M$ (wt AcpP)), we determined those parameters as well. The final concentrations of all reagents to determine $K_M$ (AcpP) and $K_M$ (malCoA) were: 50 mM sodium phosphate pH 6.8, 0.5 mM TCEP, 1 mM EDTA, 2 mM α-ketoglutaric acid, 0.4 mM $NAD^+$, 0.4 mM TPP, 15 mU/100 µL KDH, 0.025 mg/mL BSA, 10-250 µM *holo*-AcpP wt and 200 µM malCoA or 200 µM *holo*-AcpP wt and 1-100 µM malCoA, 0.02 nM FabD wt.

The data obtained by the KDH assay were fitted to the function:

$$v = k\mathrm{cat} \cdot [\mathrm{substrate}] / (K_M + [\mathrm{substrate}]) \qquad \text{eq.1}$$

The corresponding $k_{cat}$ and $K_M$ parameters are indicated on each graph (Figure S14).

**A11. Relative activities of FabD and AcpP mutants.** To most accurately report the effect of each mutation on FabD activity, each rate of transacylation reaction catalyzed by a FabD variant, or with AcpP variant as substrate, or both, was normalized to the reaction rate catalyzed by wt FabD with wt AcpP at the corresponding concentration of substrate for each replicate. Therefore, the relative rate of wt *holo*-AcpP malonylation catalyzed by wt FabD is set to 100 %. The final concentrations of all reagents were: 50 mM sodium phosphate pH 6.8, 0.5 mM TCEP, 1 mM EDTA, 2 mM α-ketoglutarate, 0.4 mM $NAD^+$, 0.4 mM TPP, 15 mU/100 µL KDH, 0.025 mg $mL^{-1}$ BSA, 50 (1x $K_M$) or 200 (4x $K_M$) µM AcpP (wt or mutants), 50 or 200 µM malonyl-CoA (malCoA) and 0.02 nM FabD (wt or mutants).

**A12. Preparation of C2-α-bromo-*crypto*-AcpP for analytical crosslinking efficiency assays.** *apo*-AcpP (1.5 mg/mL) was treated with CoaA (0.20 mg/mL), CoaD (0.20 mg/mL), CoaE (0.20 mg/mL), Sfp (0.10 mg/mL), and C2-α-bromo-pantetheine (**1**) (1.0 mM) in 50 mM Tris, pH 8 containing NaCl (50 mM), $MgCl_2$ (25 mM), ATP (30 mM), TCEP (1.0 mM), and TritonX (0.02%) at 37 °C until no *apo*-AcpP remained, as determined by 20% Urea-PAGE (~22 h). The resulting *crypto*-AcpP was purified by size-exclusion chromatography on a Superdex 75 pg (S75) column in 50 mM Tris, pH 7.4, 150 mM NaCl, 0.5 mM TCEP, and 10% glycerol.

**A13. Analytical crosslinking assays.** 5 µM FabD S92C (or 5 µM FabD S92C interface mutant) and 25 µM C2-α-bromo-*crypto*-AcpP were incubated in 50 mM Tris, pH 8 containing 0.5 mM TCEP for 26 h at 37 °C. Negative control reactions were performed simultaneously with FabD S92C (or FabD S92C interface mutant) in the absence of AcpP. Positive control reactions were performed with 5 µM FabF and 25 µM C2-α-bromo-*crypto*-AcpP. Reactions were quenched with 5x SDS loading buffer, heated at 95 °C for 5 min, and subjected to 12% SDS-PAGE, 160 V, 1 h. Gels were stained with Coomassie Brilliant Blue and scanned at 600 dpi using and EPSON Perfection V19 scanner. Coomassie staining was measured using Image Studio Lite Version 5.2 (Li-Cor Biosciences). Crosslinking efficiencies were calculated by dividing the intensity of the AcpP-FabD crosslinked band by the sum of the intensities of the AcpP-FabD crosslinked and FabD bands. Crosslinking efficiencies of each FabD S92C interface mutant were normalized to the crosslinking efficiency of FabD S92C.

While crosslinking results indicate that nearly all of the interactions identified in the AcpP-FabD structure are important for complex formation (Figure S9c), the kinetic results demonstrate that the AcpP-FabD interface can tolerate variations. We believe this difference can be explained by the relative rates of these two different reactions. The reported $k_{cat}$ values of FabD for *holo*-ACP transacylation range between 1500 and 2500 $s^{-1}$ (Figure S14),[17,18] while the crosslinking reaction between C2-α-bromo-*crypto*-AcpP and FabD S92C requires several hours to form appreciable amounts of product (Figure S28). We therefore conclude that the formation of crosslinked complex via $S_N2$ displacement of bromine by the non-natural Cys92 residue relies more upon longer-lived PPIs than the native transacylation reaction.

**A14. Time-dependent crosslinking.** 10 µM FabD S92C and 50 µM C2-α-bromo-*crypto*-AcpP were incubated in PBS, pH 7.0 for 1–48 h at 37 °C. A control was performed by incubating 10 µM FabD S92C with 50 µM *apo*-AcpP for 48 h. Reactions were quenched with 5x SDS loading buffer, heated at 95 °C for 5 min, and subjected to 12% SDS-PAGE, 160 V, 1 h. Gels were visualized with Coomassie Brilliant Blue and scanned

### B. Synthetic Protocols

**B1. General synthetic methods.** Chemical reagents were purchased from Acros, Fluka, Sigma-Aldrich, or TCI. Deuterated NMR solvents were purchased from Cambridge Isotope Laboratories. When necessary, reactions were conducted with vigorously dried anhydrous solvents that were obtained by passing through a solvent column exposed of activated A2 alumina. Air and moisture-sensitive reactions were performed under positive pressure of argon in flame-dried glassware sealed with septa and stirred with Teflon coated stir bars using an IKAMAG TCT-basic mechanical stirrer (IKA GmbH). Analytical Thin Layer Chromatograpy (TLC) was performed on Silica Gel 60 F254 precoated glass plates (EM Sciences). Visualization was achieved with UV light and/or appropriate stain ($I_2$ on $SiO_2$, $KMnO_4$, bromocresol green, dinitrophenylhydrazine, ninhydrin, or ceric ammonium molybdate). Flash column chromatography was carried out with Geduran Silica Gel 60 (40–63 mesh) from EM Biosciences. Yield and characterization data correspond to isolated, chromatographically, and spectroscopically homogeneous materials. $^{1}$H NMR spectra were recorded on Varian Mercury 400, Varian Mercury Plus 400, or JEOL ECA500 spectrometers. $^{13}$C NMR spectra were recorded at 100 MHz on Varian Mercury 400 or Varian Mercury Plus 400 spectrometers, or at 125 MHz on a JEOL ECA500 spectrometer. Chemical shifts for $^{1}$H NMR and $^{13}$C NMR analyses were referenced to the reported values of Gottlieb[19] using the signal from the residual solvent for $^{1}$H spectra, or to the $^{13}$C signal from the deuterated solvent. Chemical shift δ values for the $^{1}$H and $^{13}$C spectra are reported in parts per millions (ppm) relative to these referenced values, and multiplicities are abbreviated as s=singlet, d=doublet, t=triplet, q-quartet, m=multiplet, b=broad. All $^{13}$C NMR spectra were recorded with complete proton decoupling. FID files were processed using MestreNova 10.0 (MestreLab Research). Electrospray ionization (ESI) mass spectrometric analyses were preformed using a ThermoFinnigan LCQ Deca spectrometer. Spectral data and procedures are provided for all new compounds and copies of spectra have been provided.

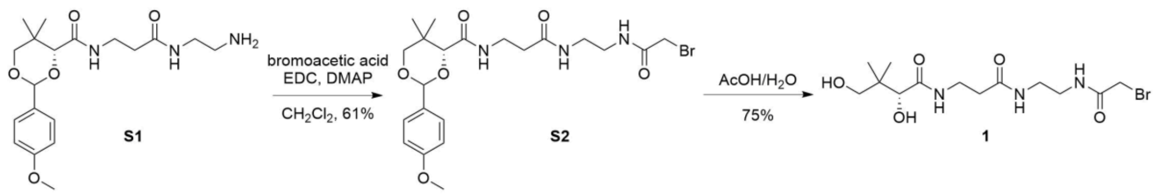

**Scheme S1. Synthesis of C2-α-bromo-pantetheineamide (1).** EDC=1-Ethyl-3-(3-dimethylaminopropyl)carbodiimide); DMAP=4-(dimethylamino)-pyridine; AcOH=acetic acid.

**(4*R*)-*N*-(3-((2-aminoethyl)amino)-3-oxopropyl)-2-(4-methoxyphenyl)-5,5-dimethyl-1,3-dioxane-4-carboxamide (S2).** In a 10 mL pear-shaped flask, (4*R*)-*N*-(3-((2-aminoethyl)amino)-3-oxopropyl)-2-(4-methoxyphenyl)-5,5-dimethyl-1,3-dioxane-4-carboxamide **S1**[20] (102.0 mg, 0.2688 mmol, 1.0 equiv.), bromoacetic acid (58.6 mg, 0.422 mmol, 1.6 equiv.), and 3 mL $CH_2Cl_2$ were added. To the solution was added EDC•HCl (81.6 mg, 0.426 mmol, 1.6 equiv.) and DMAP (9.7 mg, 0.0794 mmol, 0.30 equiv.). After 21 h, the reaction was dissolved in 100 mL $CH_2Cl_2$ and washed with deionized water (10 mL). The organic phase was dried over $MgSO_4$, filtered, and concentrated by rotary evaporation. Purification by silica flash chromatography (39:1 □ 9:1 $CH_2Cl_2$/MeOH) afforded **S2** (81.6 mg, 60.7%) as a white solid.

**TLC:** $R_f$ 0.14 (19:1 $CH_2Cl_2$/MeOH). **$^1$H-NMR** (500 MHz, $CDCl_3$): δ 7.40 (d, *J* = 10.8 Hz, 2H), 7.06–6.99 (m, 2H), 6.88 (d, *J* = 8.8 Hz, 1H), 5.43 (s, 1H), 4.04 (s, 1H), 3.94 (s, 1H), 3.78 (s, 3H), 3.75 (s, 1H), 3.70–3.59 (m, 2H), 3.49 (s, 2H), 3.32 (s, 4H), 2.46–2.34 (m, 2H), 1.05 (d, *J* = 12.2 Hz, 6H).**$^{13}$C-NMR** (125 MHz, $CDCl_3$): δ 172.17, 169.79, 167.17, 160.33, 130.17, 127.63, 113.81, 101.42, 83.91, 78.48, 55.44, 42.59, 40.45, 39.23, 36.11, 35.16, 33.15, 28.92, 21.94, 19.22. **HR-ESI-MS** *m/z* calcd. $[C_{21}H_{30}BrN_3O_6Na]^+$: 522.1210, found 522.1206.

**(R)-N-(3-((2-(2-bromoacetamido)ethyl)amino)-3-oxopropyl)-2,4-dihydroxy-3,3-dimethylbutanamide (C2-α-bromo-pantetheineamide, 1).** In a 20 mL vial, **S2** (81.6 mg, 0.163 mmol, 1.0 equiv.) and1.0 mL 4:1 AcOH/$H_2O$ were added. After 7 h, the mixture was concentrated by rotary evaporation, then azeotroped from cyclohexane (5x10 mL) and benzene (3x10 mL). Purification by silica flash chromatography (19:1 □ 3:1 $CH_2Cl_2$/MeOH) afforded **1** (47.0 mg, 75.4%) as a clear oil. Analytical data were consistent with previous reports.[21]

**TLC:** $R_f$ 0.22 (9:1 $CH_2Cl_2$/MeOH). **$^1$H-NMR** (400 MHz, $CD_3OD$): δ 4.06 (s, 2H), 3.87 (d, *J* = 16.4 Hz, 2H), 3.54–3.42 (m, 4H), 3.42–3.28 (m, 5H), 2.42 (t, *J*=6.6 Hz, 2H), 0.91 (s, 6H).**$^{13}$C-NMR** (100 MHz, $CD_3OD$): δ 174.87, 173.09, 168.50, 76.20, 69.11, 42.04, 39.37, 39.19, 38.60, 35.44, 35.23, 20.28, 19.68. **HR-ESI-MS** *m/z* calcd. $[C_{13}H_{24}BrN_3O_5Na]^+$: 404.0792, found 404.0793.

### C. Molecular Dynamics Simulations Protocols

**C1. Structure preparation.** As shown in Table 3, a total of 17 systems were subjected to computer simulations. For simulation work, the coordinates of the wildtype and mutant *apo*-FabD, malonyl-CoA•FabD, and malonyl-AcpP•FabD structures were generated using x-ray crystal structures of *apo*-FabD (PDB ID: 1MLA, 2G1H)[22,23] malonyl-CoA•FabD (PDB ID: 2G2Z)[23], and the crosslinked AcpP-FabD (reported herein), respectively. Mutants of the enzymes and enzyme complexes were generated using Pymol v.2.3[24] (https://pymol.org/2/support.html) and Schrodinger's Protein Preparation Wizard[24–26] (https://www.schrodinger.com/protein-preparation-wizard[27,28]) and https://www.schrodinger.com/prime were used to add missing C-, N-terminal residues and missing side chains not resolvable from the experimental density, and hydrogen atoms were added all heavy atoms  to cap all open valences. To predict the protonation states of the titratable residues in each structure assuming a pH of 7.4, and to optimize the orientation, all waters resolved crystallographically. Note that in the preparation of all protein structures the active site S92C mutation of the FabD was "reversed" in order to restore the active site serine found in the wildtype enzyme. Histidine protonation states were inspected by hand.

**C2. Simulation preparation.**

In an analogous manner, TLEAP (https://ambermd.org/CiteAmber.php) were used to generate AMBER (ff14SB/GAFF2) topology and the parameters files for the simulation cell of all 17 systems of interest. Parameterization of the malonyl-CoA and malonyl-AcpP's nonstandard residue was performed using ANTECHAMBER and GAFF2 force field (https://ambermd.org/CiteAmber.php).[29,30] Subsequently topology and parameter files were prepared using TLEAP. TLEAP was used to prepare the simulation cell in a manner analogous to that of the CHARMM-GUI approach. All simulation cells were prepared using TIP3P water molecules[31] to generate isometric cell to ensure that the cell walls are 10 Å away from the closet portion of the coordinates derived from experimental data in either data. The restrained electrostatic potential (RESP) method[32] was used to determined partial atomic charges for the ANTECHAMBER parameterization of the malonyl-containing nonstandard residue.[29] The potential was computed at the HF/6-31G(d) level of theory using Gaussian 09 (https://gaussian.com/g09citation/).

**C3. Simulation methodology.** Simulations were performed using GPU-accelerated Amber 2016 and Amber 2018 (https://ambermd.org/CiteAmber.php).[33,34] A 2 fs time-step was utilized via the

SHAKE algorithm, which constrains all nonpolar bonds involving hydrogen atoms.[35] Long-range electrostatic interactions were treated using the Particle Mesh Ewald (PME) method with a 10 Å cutoff for all non-bonded interactions.[36] Both solvated protein complexes were energy minimized in a two-step fashion. In a first step, solvent molecules and counterions were allowed to relax, while all protein atoms were restrained using a harmonic potential (k = 500 kcal $mol^{-1}$ $Å^{-2}$). This geometry optimization was followed by an unrestrained energy minimization of the entire system. The thermal energy available at a physiological temperature of 310 K was slowly added to each system over the course of a 2 ns NVT ensemble simulation. The solvated complexes were then subjected to unbiased isobaric-isothermal (NPT) simulations for 10 ns in order to equilibrate the heated structures. Three independent 512 ns production MD (NPT ensemble) of each system were performed with different initial velocities. For both NVT and NPT simulations, the Langevin thermostat ($\lambda$ = 5.0 $ps^{-1}$) was used to maintain temperature control.[37–39] Pressure regulation in NPT simulations (target pressure of 1 atm) was achieved by isotropic position scaling of the simulation cell volume using a Berendsen barostat.[40] Coordinate data was written to disk every 10 ps.

**C4. Analysis and visualization of simulation data:** Analysis was performed using CPPTRAJ,[41] PYTRAJ, a Python front-end for the CPPTRAJ analysis code (https://amber-md.github.io/pytraj/latest/overview.html#citations). Trajectories were visualized using NGLview[42–44] and Pymol v2.3 (https://gaussian.com/g09citation/). All data was plotted using matplotlib library of Python.

The process of modifying the experimental structure through structure and simulation preparation modifies its geometry such that there is a modest difference between the experiment and initial coordinates (for simulation). Furthermore, because the crosslinker used to trap the experimental AcpP-FabD complex was replaced with malonyl moiety in the structures prepared for simulation, some analysis including native contact analyses were performed using the initial coordinates instead of the experimental ones.

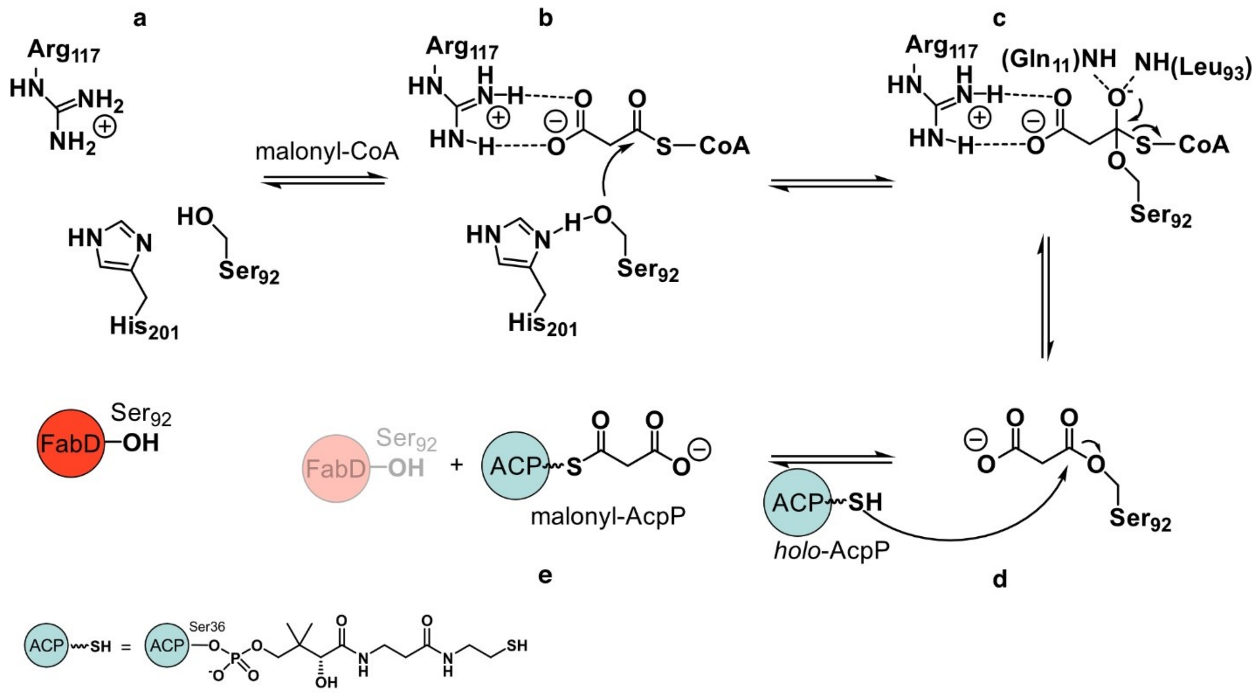

**Figure S1. Reaction catalyzed by FabD** (adapted from references[23,45,46]). **a**) FabD active site residues. **b**) FabD-catalyzed incorporation of malonyl-CoA occurs *via* a ping–pong bi–bi mechanism involving the malonylation of FabD catalytic serine (Ser92) in the first step. **c**) The tetrahedral intermediate is stabilized by an oxyanion hole formed with the backbone amides from Gln11 and Leu93. **d**) The acyl-FabD intermediate is then subject to a nucleophilic attack by the thiol of the phosphopantetheine arm of *holo*-AcpP, **e**) which results in the formation of malonyl-AcpP.

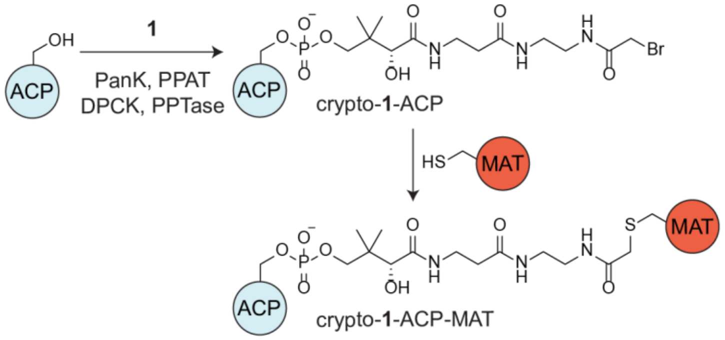

**Figure S2. Chemoenzymatic modification of *apo*-AcpP with C2-α-bromo-pantetheine (1) and subsequent FabD S92C crosslinking.** *apo*-ACP is modified with 1 to form *crypto*-1-ACP, followed by nucleophilic attack by the active site cysteine from the MAT to form crosslinked ACP-MAT complex. In situ conversion of pantetheine analog 1 is converted into a coenzyme A analog using pantothenate kinase (PanK), 4′-phosphopantetheine adenyltransferase (PPAT), and dephospho-coenzymeA kinase (DPCK), followed by *B. subtilis* phosphopantetheinyl transferase (PPTase)-catalyzed loading of phosphopantetheine analog onto *apo*-ACP.

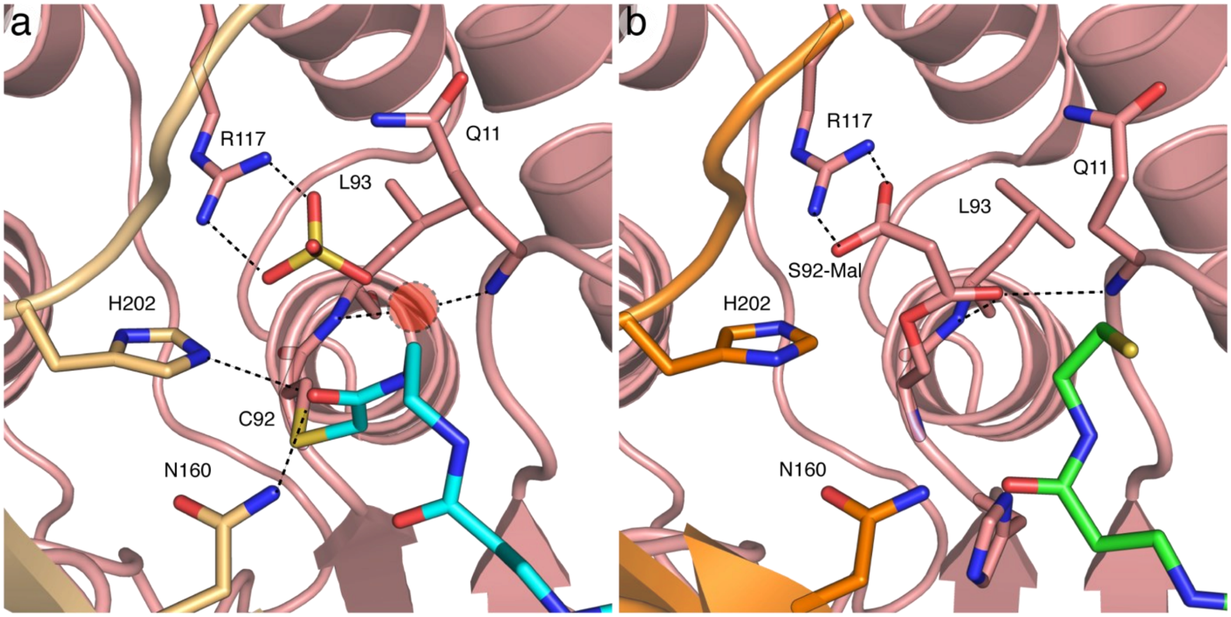

**Figure S3. Comparison of AcpP-FabD and malonyl-CoA•FabD active site interactions. a)** Active site of FabD from AcpP-FabD structure highlighting interactions between the acetamide portion of the PPant crosslinking probe (carbon colored cyan) and the bound sulfate ion in the active site. The acetamide carbonyl group is found rotated away from the oxyanion hole and instead engages in hydrogen-bonding interactions with His202 and Asn160. The oxyanion hole, formed by backbone amides of Gln11 and Leu93, is shown as red transparent circle with a dotted outline. Residues from the FL subdomain have light-orange colored carbon atoms and residues from the ABH subdomain have salmon colored carbon atoms. **b)** Active site of FabD from malonyl-CoA•FabD showing the carbonyl group of bound malonyl-Ser92 coordinated in the active site oxyanion hole and bidentate salt-bridge interaction between the carboxylate of malonyl-Ser92 with Arg117. The bound hydrolyzed CoA moiety (carbon colored green) is also shown in the active site. Important interactions within a hydrogen bonding distance cutoff of 3.5 Å are shown by dotted lines. Residues from the FL subdomain have orange colored carbon atoms and residues from the ABH subdomain have salmon colored carbon atoms.

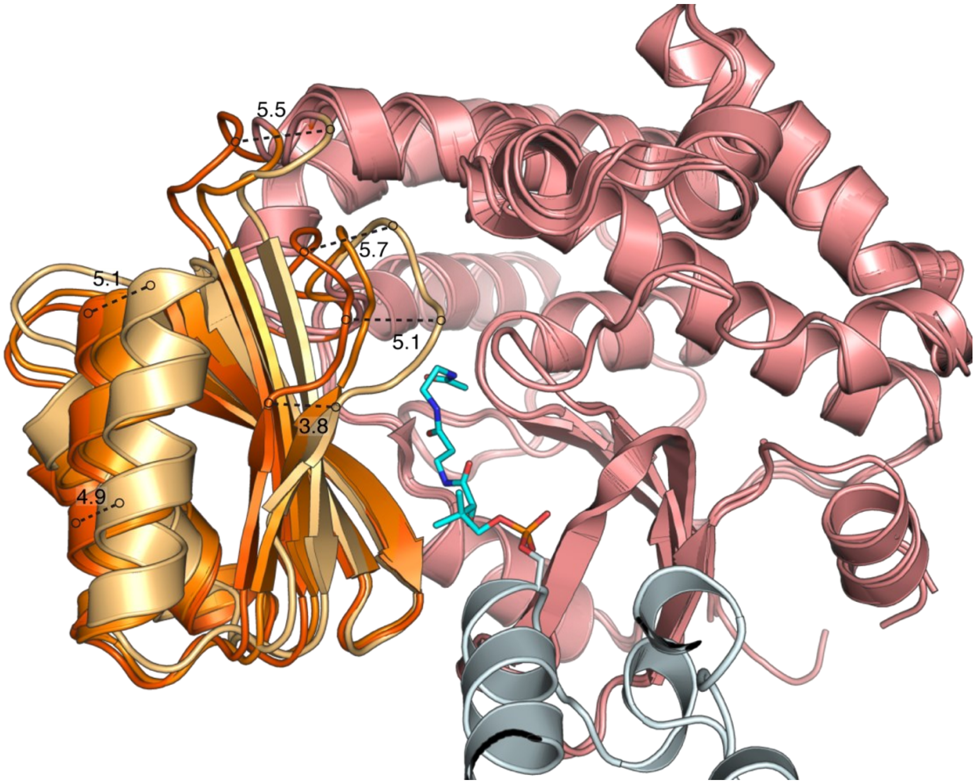

**Figure S4. Measurement of FL subdomain rigid body motions.** Measurements of the rigid body domain motions of the FL subdomain with respect to the ABH subdomain in the apo-FabD (PDB ID: 1MLA, dark orange) and AcpP-FabD (PDB ID: 6U0J, light orange) structures (distance given in Å). The malonyl-Coa-FabD complex (PDB ID: 2G2Z, orange) is also displayed for consistency with the main text figure.

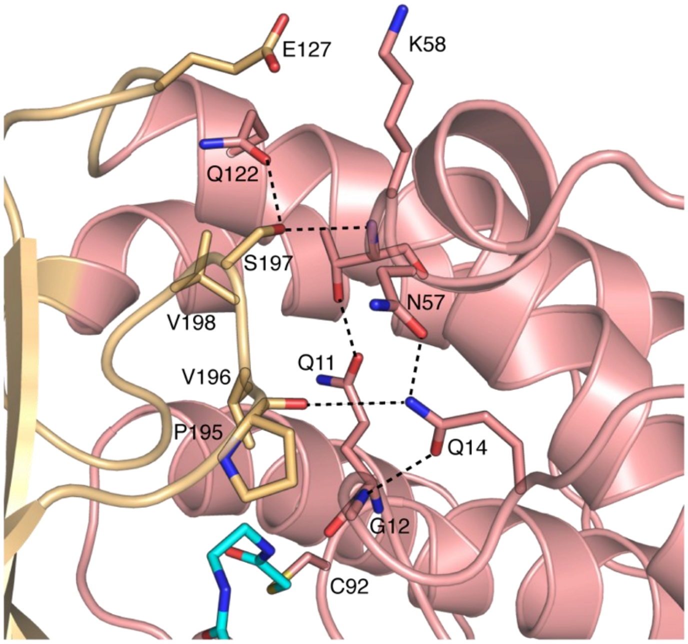

**Figure S5. Interaction networks formed between FL and ABH subdomains in AcpP-FabD structure.** Rigid body subdomain motion of the FL subdomain results in new contacts formed between the ABH and FL subdomain in the AcpP-FabD structure. Residues along the β4-loop form a new network of contacts and important interactions within a hydrogen bonding distance cutoff of 3.5 Å are shown by dotted lines. Residues from the FL subdomain have light-orange colored carbon atoms and residues from the ABH subdomain have salmon colored carbon atoms. The PPant arm is shown crosslinked to the active site Cys92 residue and carbons are colored cyan.

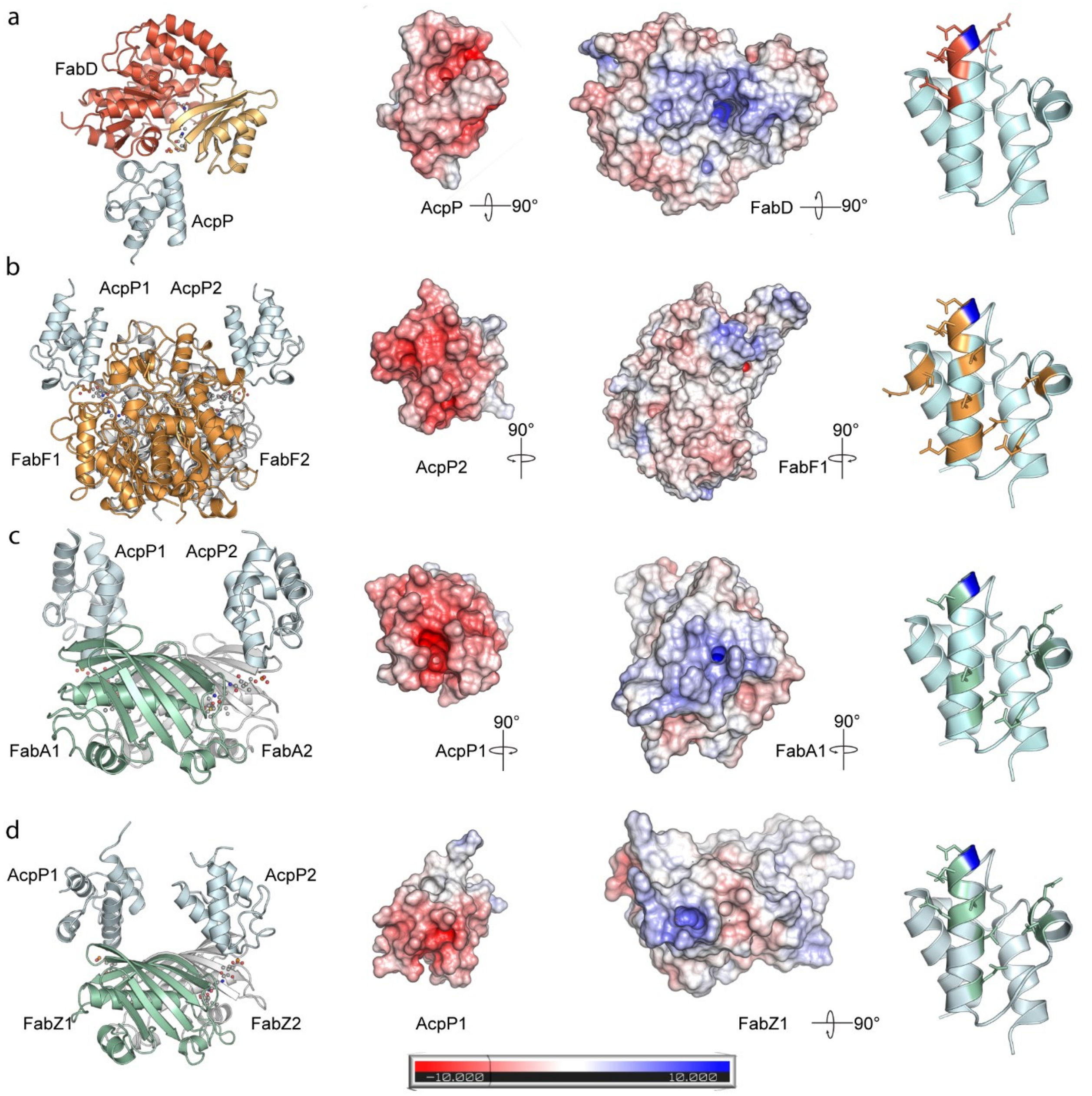

**Figure S6. AcpP Interactome.** Structural comparison of the *E. coli* (**a**) AcpP-FabD (PDB ID: 6U0J), (**b**) AcpP-FabF (PDB ID: 6OKG) (**c**) AcpP-FabA (PDB ID: 4KEH), and (**d**) AcpP-FabZ dimer (PDB ID: 6N3P) crosslinked complexes, depicting the overall structural architecture (left), electrostatic potential (ESP) maps of the partner protein and AcpP monomers (middle), and *apo*-AcpP structure (PDB: 1T8K, 1.1 Å), illustrating contacts that each partner protein makes with the AcpP (right), with the phosphopantetheinylation site (Ser36) depicted in blue. In all cases, the ESP is mapped onto the solvent-excluded surface (Connolly surface) of the partner protein and AcpP monomers using a blue to white to red color scheme spanning from –10.0 $kTe^{-1}$ to +10.0 $kTe^{-1}$. Note that AcpP-FabZ structure is a hexamer consisting of three dimers. Here, one of those dimers is depicted in panel **d**.

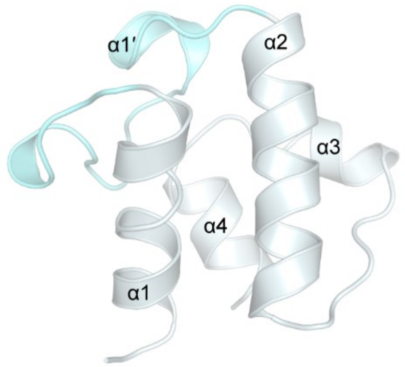

**Figure S7. Overall topology of AcpP in the AcpP-FabD structure** (PDB ID: 6U0J). AcpP forms contacts with FabD through a small helix in loop 1 (α1′) and the N-terminal region of helix II (α2). The loop 1 region is colored aquamarine.

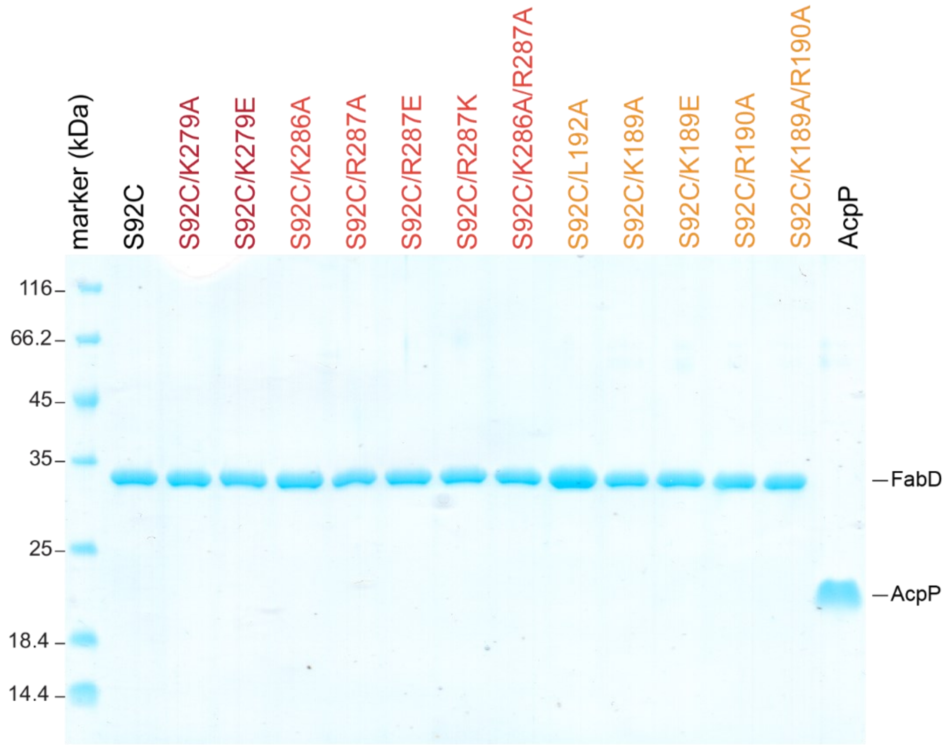

**Figure S8. Purified FabD S92C interface mutants for crosslinking validation.** Coomassie-stained 12% SDS-PAGE gel of FabD S92C interface mutants and C2-α-Br-pantetheinamide-AcpP.

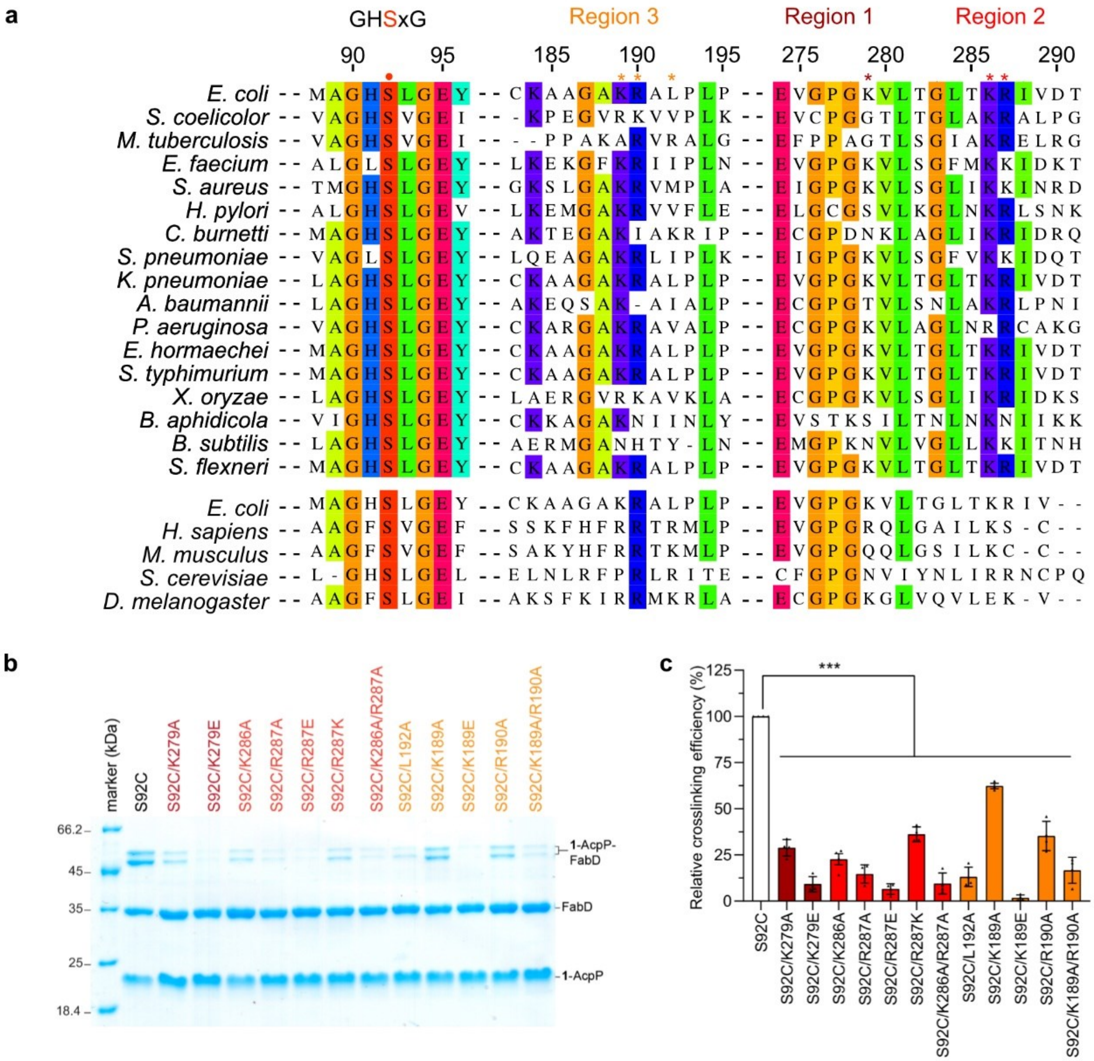

**Figure S9. Validation of FabD-AcpP interface via mutagenesis and crosslinking. a)** Sequence alignment of MATs from type II FAS (top) and type II eukaryotic mitochondrial MATs (bottom). The catalytic serine of FabD, part of the highly conserved GHSxG motif in ATs active site, is denoted by a red circle. Residues that were mutated for crosslinking and kinetics studies are denoted by asterisks. Residues are colored based on sequence identity (threshold set at 70%). **b)** SDS-PAGE analysis depicting crosslinking between 5 µM FabD S92C interface mutants and 25 µM C2-α-bromo-*crypto*-AcpP in 50 mM Tris, pH 8 containing 0.5 mM TCEP at 37 °C for 26 h. **c)** Semi-quantitative densitometric analysis of SDS-PAGE data, showcasing FabD interface residues from region 1 (crimson), region 2 (red), and region 3 (orange). The mutants were subjected to a crosslinking assay using C2-α-bromo-*crypto*-AcpP and all reported data were normalized to FabD S92C crosslinking efficiency. Data is reported as the mean ± standard deviation from 4 replicates. Statistical significance was compared using Dunnett's multiple comparison test. ***$p<0.001$.

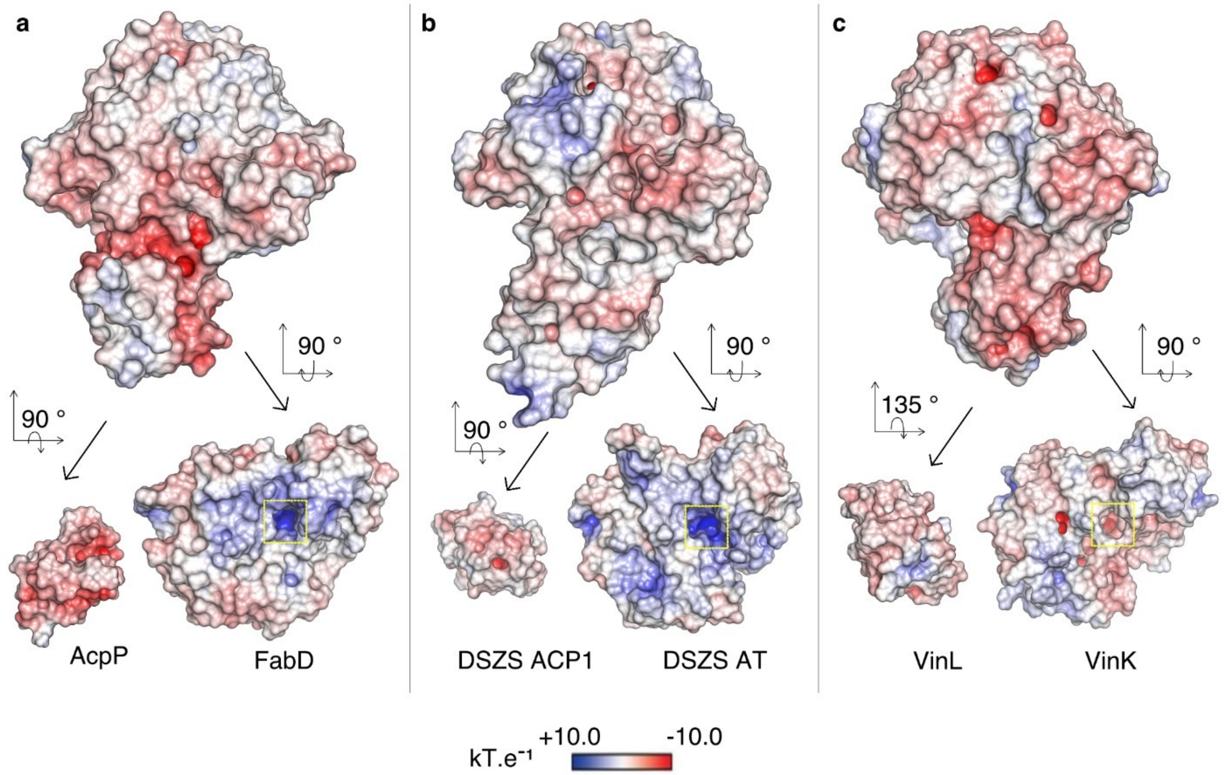

**Figure S10. ACP-AT electrostatic potentials (ESP) maps of ACP-AT complexes and respective monomers. a**) AcpP-FabD (PDB ID: 6U0J), **b**) DSZS ACP1-DSZS AT (PDB ID: 5ZK4), **c**) VinL-VinK (PDB ID: 5CZD). In all cases, the ESP is mapped onto the "Connelly" surfaces of the AT and ACP monomers using a blue to white to red color range, spanning from +10.0 kT•e$^{-1}$ to –10.0 kT•e$^{-1}$. The entrance of the AT active site is indicated by the yellow box.

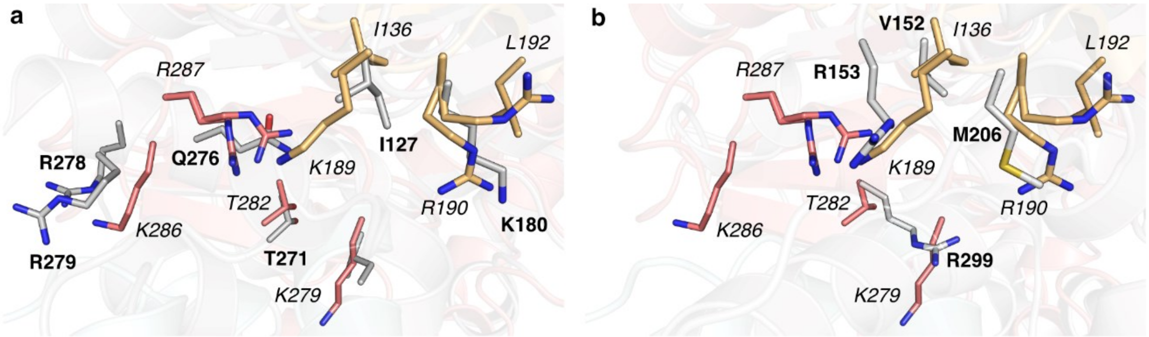

**Figure S11. Interfaces residues of *trans*-ATs compared to FabD. a)** AcpP-FabD (PDB ID: 6U0J) overlaid with DSZS ACP1-DSZS AT (PDB ID: 5ZK4). **b)** AcpP-FabD overlaid with VinL-VinK (PDB ID: 5CZD). Only AT interface residues are shown for clarity. Interface residues from the *trans*-ATs (grey) and from FabD (colored) are shown in bold and in italic, respectively.

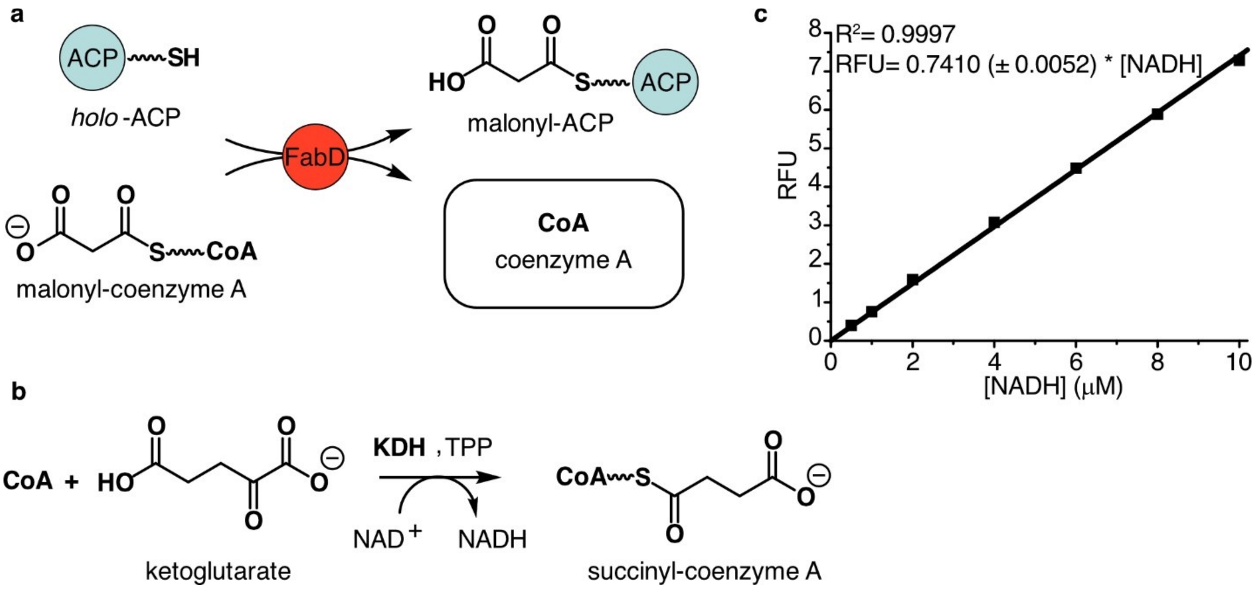

**Figure S12. FabD continuous coupled assay using α-ketoglutarate dehydrogenase (KDH).** KDH is a nicotinamide adenine dinucleotide ($NAD^+$)- and thiamine pyrophosphate- (TPP) dependent enzyme which reacts with the free coenzyme A (CoA) generated by FabD (**a**). This reaction is accompanied with a reduction of $NAD^+$ to NADH which can be monitored by fluorescence (**b**). **c)** NADH calibration curve (each point was repeated in triplicate).

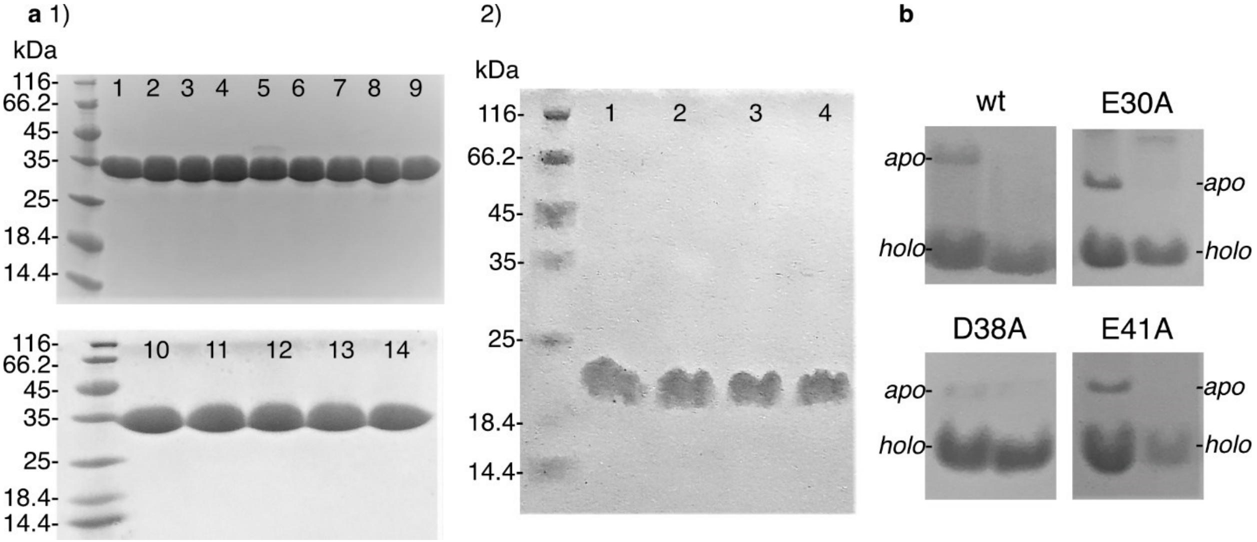

**Figure S13. a) Coomassie-stained SDS-PAGE gels** (15 % acrylamide, 170 V, 60 min) of FabD wt and mutants (**1**): FabD wt (1), K279E (2), R287A (3), R287E (4), R287K (5), S295A (6), K279A (7), K286A (8), L192A (9), K189A (10), K189E (11), R190A (12), K189A R190A (13), K286A R287A (14), and of AcpP wt and mutants (**2**): AcpP wt (1), E30A (2), D38A (3), E41A (4). **b) Coomassie-stained urea-PAGE gels** (20 % acrylamide, 170 V, 90 min) of AcpP wt and mutants. AcpP is expressed in both *apo* and *holo* form (left lane) and is fully converted to its *holo* form after reaction with Sfp (right lane).

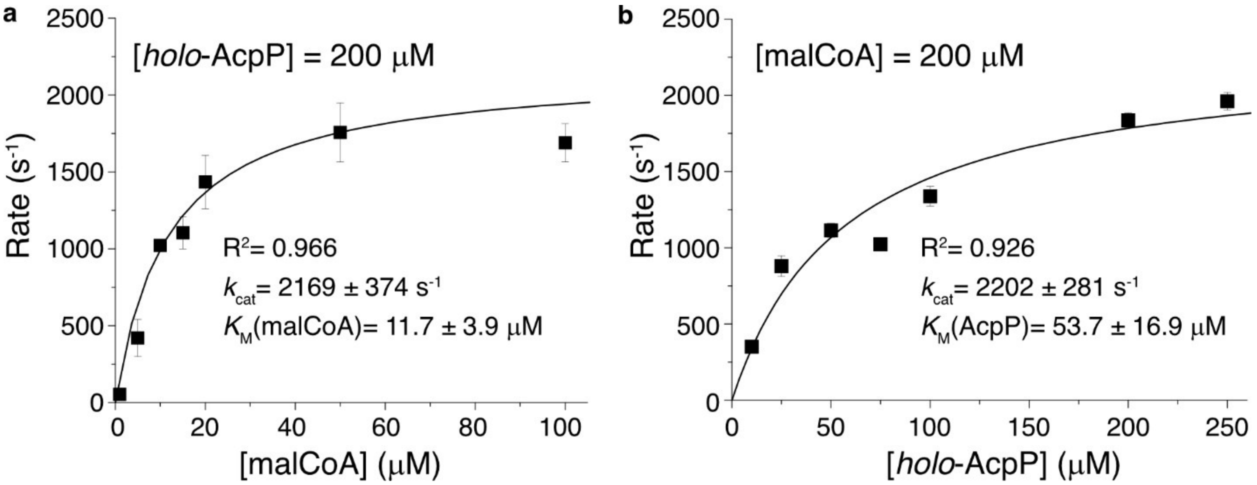

**Figure S14. Michaelis-Menten curves of FabD wt. a)** The concentration of *holo*-AcpP is kept constant (200 µM or 4 x $K_M$(AcpP)) to determine $K_M$(malCoA). **b)** The concentration of malCoA is kept constant (200 µM or > 4x $K_M$(malCoA)) to determine $K_M$(AcpP). Each reaction was run in biological triplicate and the error bars indicate the standard deviation on the mean of the three measurements.

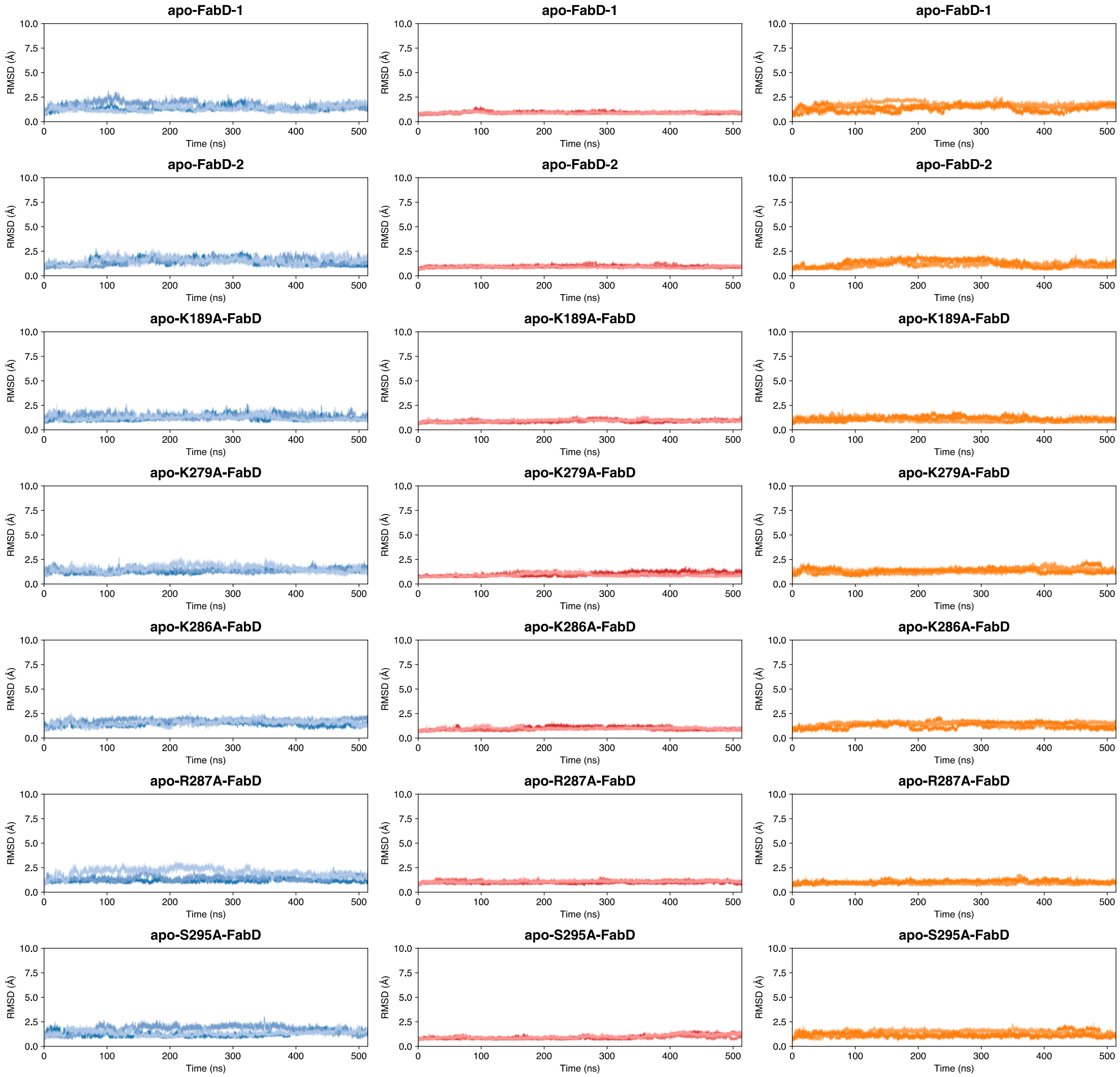

**Figure S15. $C_\alpha$ root mean square analysis (RMSD) in Ångstroms (Å) of the simulations of wildtype (wt) *apo*-FabD and five FabD mutants**. Analysis was performed on 3 independent 524 ns molecular dynamics (MD) simulations. Each plot is labeled to indicate the FabD system analyzed. The RMSDs of the entire FabD domain (left, blue curves), the ABH (large) subdomain (center, red curves), and FL (small) subdomain (right, orange curves) of FabD simulations.

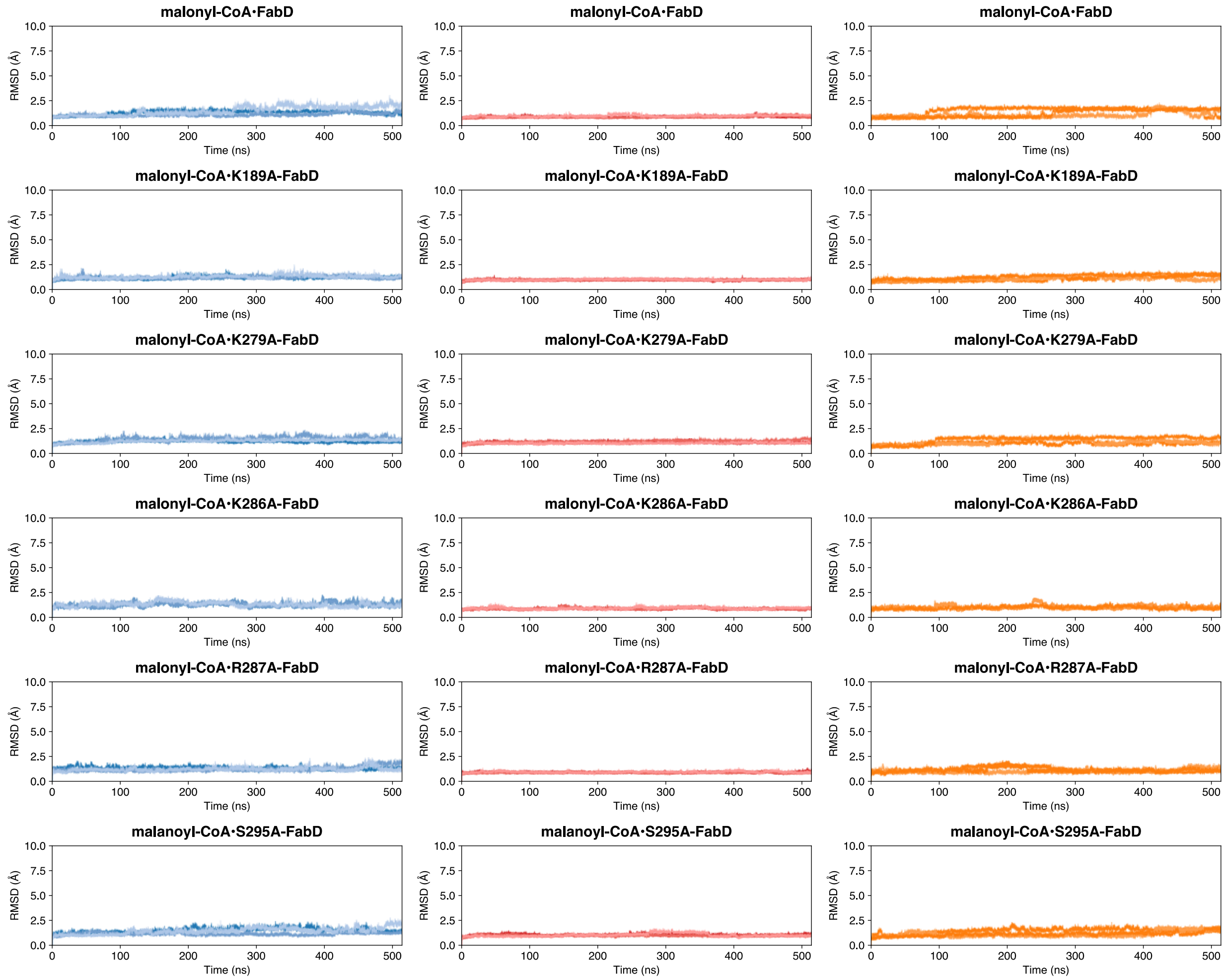

**Figure S16. $C_\alpha$ root mean square analysis (RMSD) in Ångstroms (Å) of the simulations of wildtype (wt) FabD and *apo*-FabD bound with malonyl-CoA** performed using ff14SB/GAFF2 force field parameter. Analysis was performed on 3 independent 524 ns molecular dynamics (MD) simulations. Each plot is labeled to indicate the FabD system analyzed. The RMSDs of the entire FabD domain (left, blue curves), the ABH (large) subdomain (center, red curves), and FL (small) subdomain (right, orange curves) of FabD. Analysis was performed on 3 independent 524 ns molecular dynamics (MD) simulations. Each plot is labeled to indicate the FabD system analyzed and show the change in protein RMSD over the course of each 524 ns simulation (shown in a blue, red, or orange hue).

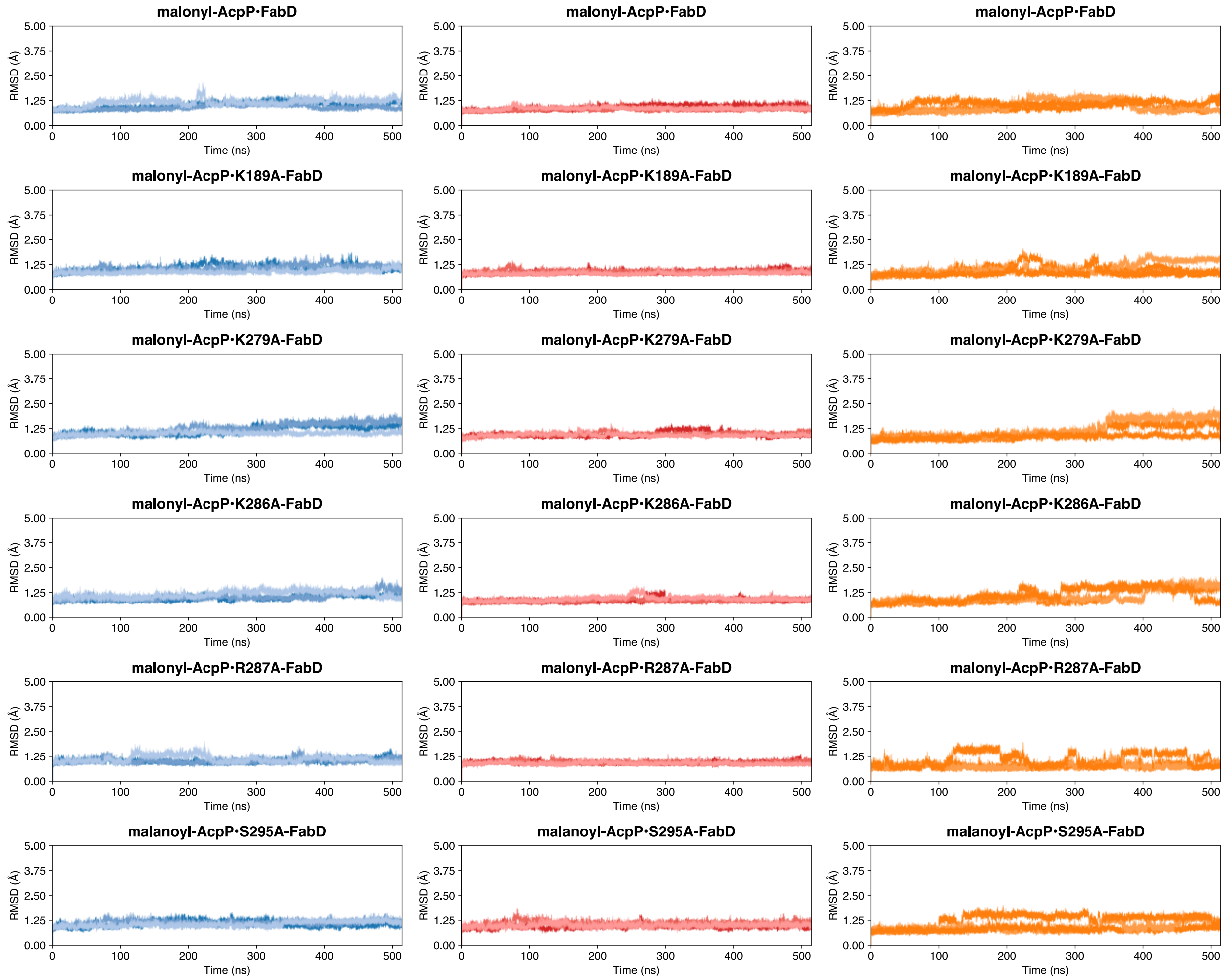

**Figure S17. C$_\alpha$ root mean square analysis (RMSD) in Ångstroms (Å) of the simulations of malonyl-AcpP-bound wildtype (wt) FabD and *apo*-FabD. Analysis was performed on 3 independent 524 ns molecular dynamics (MD) simulations**. Each plot is labeled to indicate the FabD system analyzed. The RMSDs of the entire FabD domain (left, blue curves), the ABH (large) subdomain (center, red curves), and FL (small) subdomain (right, orange curves) of FabD. Analysis was performed on 3 independent 524 ns molecular dynamics (MD) simulations. Each plot is labeled to indicate the FabD system analyzed and show the change in protein RMSD over the course of each 524 ns simulation (shown in a blue, red, or orange hue).

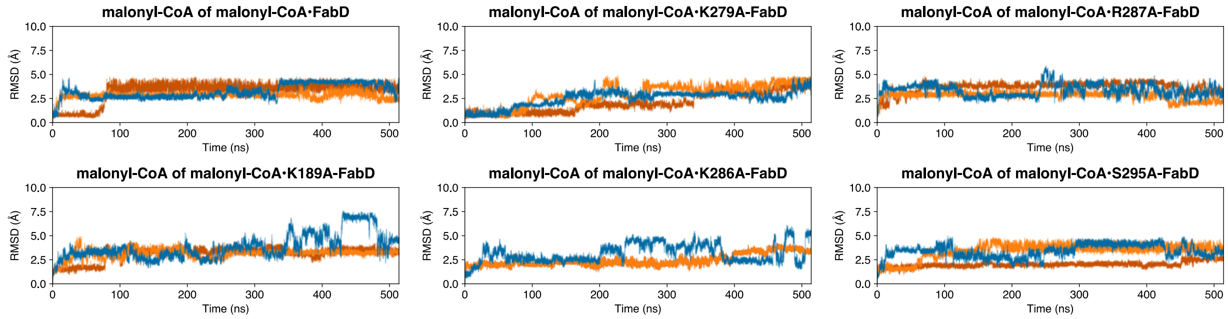

**Figure S18. Heavy atom root mean square deviation analysis (RMSD) in Ångstroms (Å) of the malonyl-CoA substrate of the wt and mutant malonyl-CoA•FabD complexes**. Analysis was performed on 3 independent 524 ns molecular dynamics (MD) simulations. Each plot is labeled to indicate the FabD system analyzed. The nonstandard MPS residues includes the conserved Ser36 of AcpP, the phosphopantetheine prosthetic arm, and the ligated malonyl moiety.

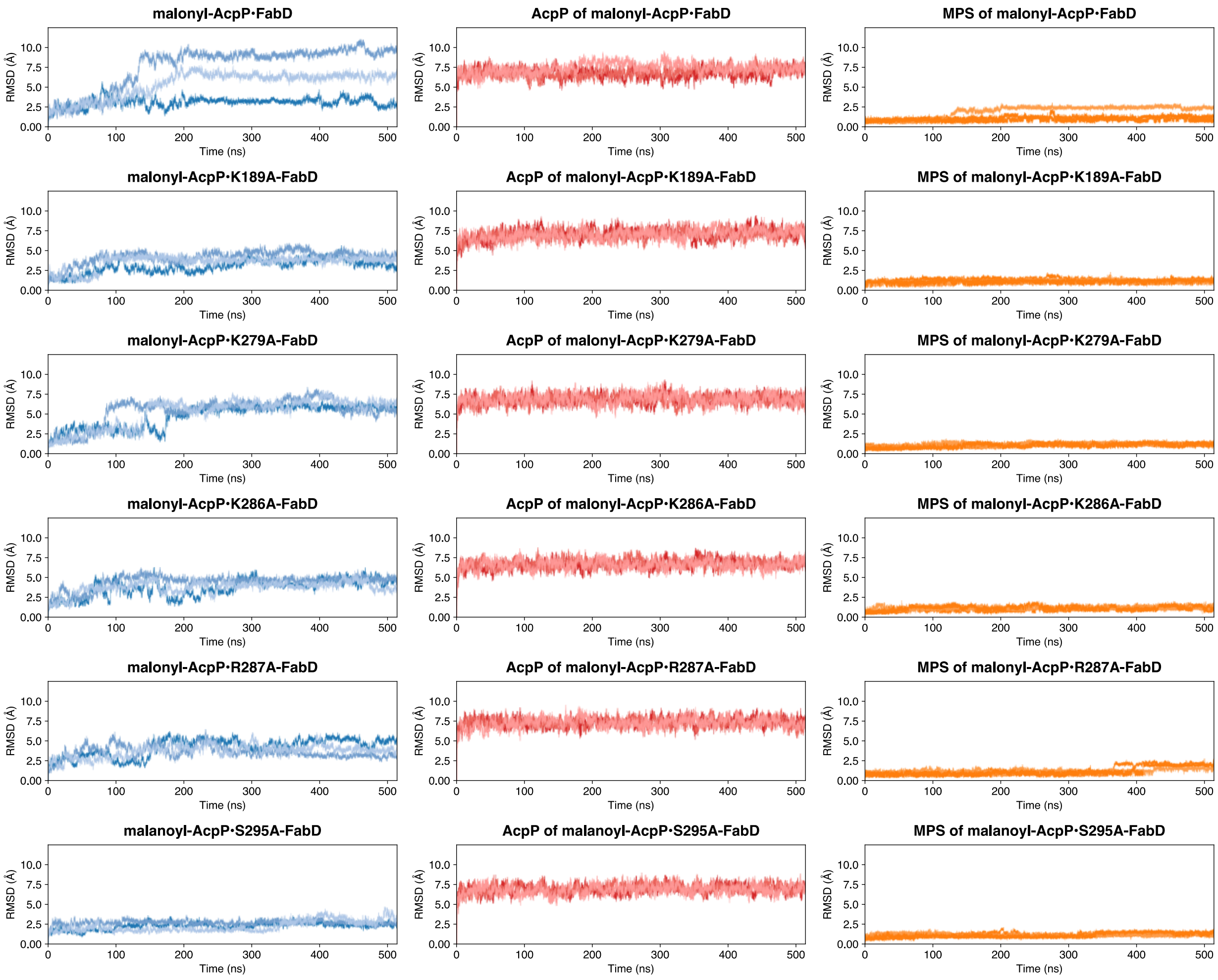

**Figure S19. Heavy atom root mean square analysis (RMSD) in Ångstroms (Å) of the wt and mutant malonyl-FabD•FabD complexes**. Analysis was performed on 3 independent 524 ns molecular dynamics (MD) simulations. Each plot is labeled to indicate the malonyl-AcpP•FabD system analyzed. The RMSDs of the entire AcpP•FabD complex (left, blue curves), the AcpP monomer (center, red curves), and nonstandard residue (right, orange curves). Each plot is labeled to indicate the FabD system analyzed and show the change in protein RMSD over the course of each 524 ns simulation (shown in a blue, red, or orange hue). The nonstandard MPS residues includes the conserved Ser36 of AcpP, the phosphopantetheine prosthetic arm, and the ligated malonyl moiety. Simulation data was superimposed over entire AcpP•FabD complex.

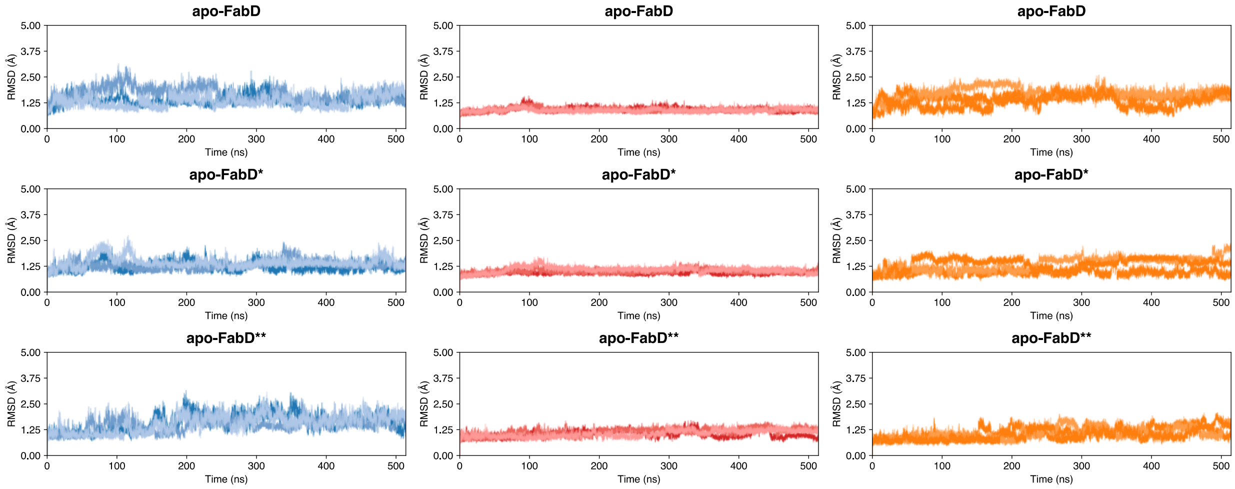

**Figure S20. $C_\alpha$ root mean square analysis (RMSD) in Ångstroms (Å) of the simulations of wildtype (wt) *apo*-FabD*(apo-FabD derived from malonyl-AcpP•FabD) and *apo*-FabD**(apo-FabD derived from malonyl-CoA•FabD)**. Analysis was performed on 3 independent 524 ns molecular dynamics (MD) simulations. Each plot is labeled to indicate the FabD system analyzed. The RMSDs of the entire FabD domain (left, blue curves), the ABH (large) subdomain (center, red curves), and FL (small) subdomain (right, orange curves) of FabD.

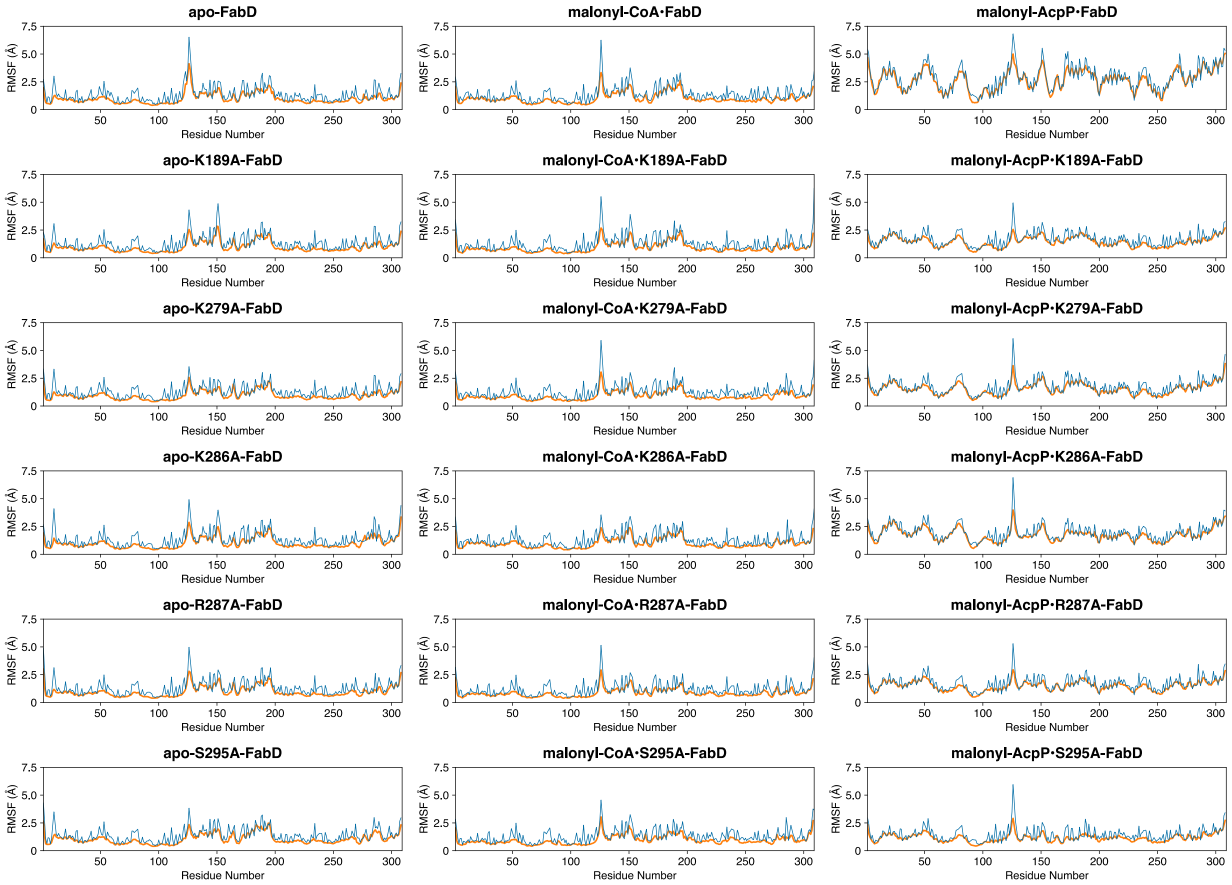

**Figure S21**. **Root mean square fluctuations (RMSFs) of each residue of FabD of the variants of the *apo*-FabD, malonyl-CoA•FabD, and malonyl-AcpP•FabD systems subjected to MD simulation**. The blue curve shows the RMS fluctuations of the backbone – measured using backbone heavy (non-hydrogenic) atoms – of each residue, whereas the orange curve shows the RMS fluctuation of sidechains measured using each residue's sidechain heavy (non-hydrogen) atoms. Coordinate data was superimposed overall all backbone atoms.

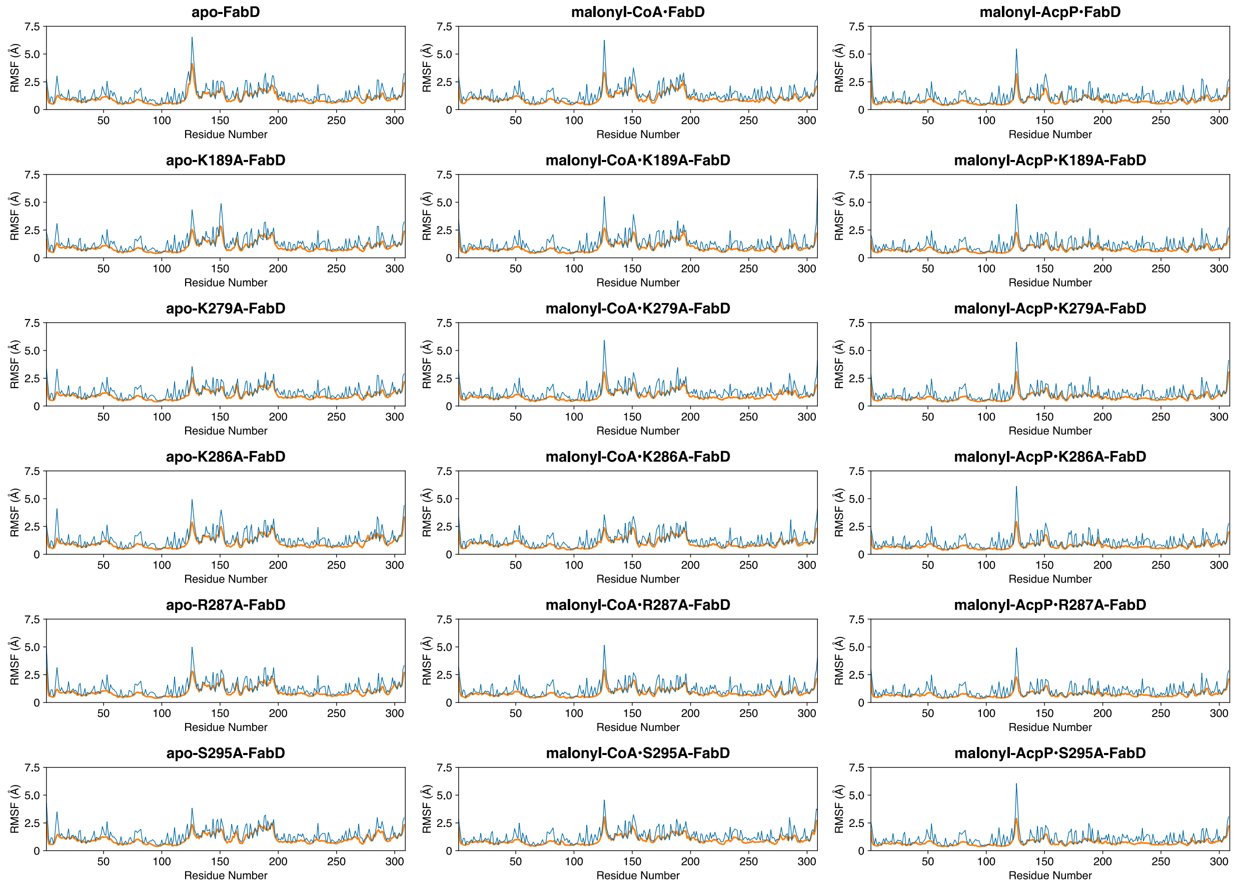

**Figure S22**. **Root mean square fluctuations (RMSFs) of each residue of FabD of the variants of the *apo*-FabD, malonyl-CoA•FabD, and malonyl-AcpP•FabD systems subjected to MD simulation**. The blue curve shows the RMS fluctuations of the backbone – measured using backbone heavy (non-hydrogenic) atoms – of each residue, whereas the orange curve shows the RMS fluctuation of sidechains measured using each residue's sidechain heavy (non-hydrogen) atoms. Coordinate data was superimposed overall backbone atoms of the FabD components of each system.

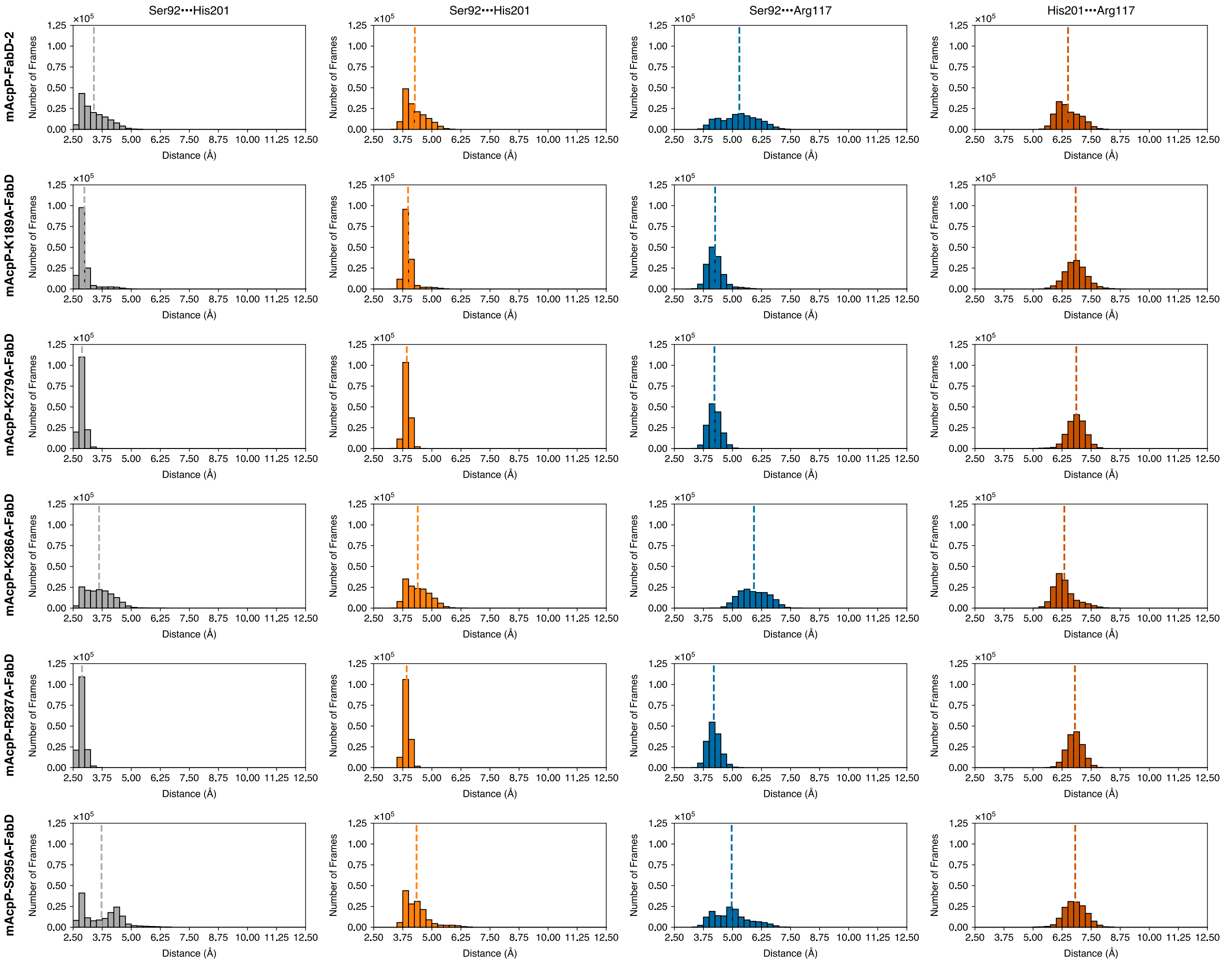

**Figure S23. Distribution of distances between key active site residues sampled throughout the course of MD simulations of malonyl-AcpP•FabD variants.** From left to right the distribution of distances between Ser92 and His201 (measured in two different ways in the 1$^{st}$ and 2$^{nd}$ columns), Ser92 and Arg117, and His201 and Arg 117 (from left to right). The distances between Ser92 and His201 were measured as distance between Oγ of Ser92 and His92's Nε2 (1$^{st}$ column) or as the distance between Oγ of Ser and the center of geometry of His201's imidazole ring. The Ser92-Arg117 is measured between Oγ of Ser and the center of geometry of Arg117's guanidinium moiety. Lastly, the distance between His201 and Arg117 is determined the centers of geometry of the imidazole and guanidinium moiety of His201 and Argy117, respectively. The dotted vertical line indicates the average distance. Distances are measured in Å. 1D histograms was generated using a bin width of 0.25 Å.

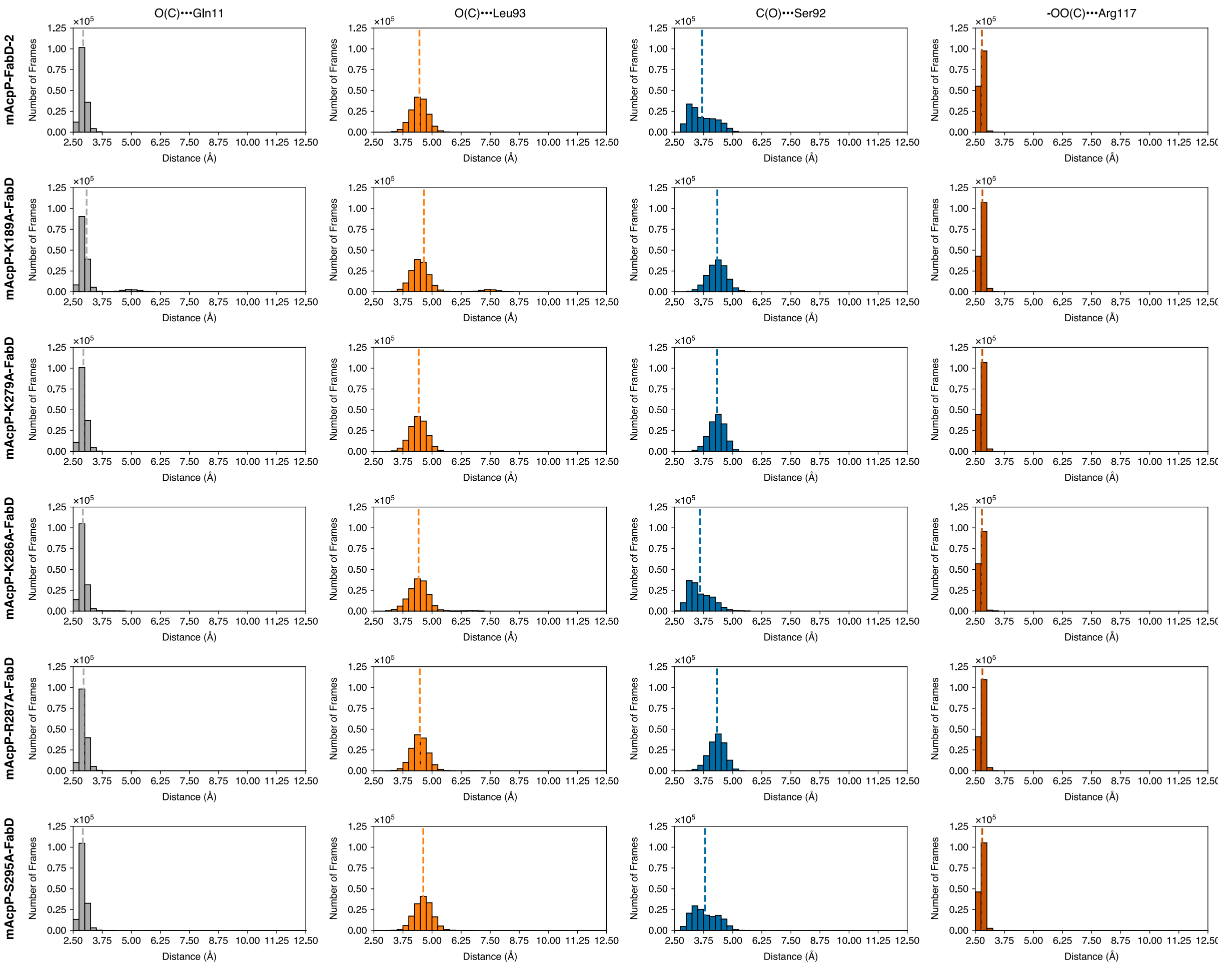

**Figure S24. Distribution of distances between key active site residues and malonyl portion of malonyl-AcpP sampled throughout the course of MD simulations of malonyl-AcpP•FabD variants.** From left to right, the distribution of distances between the backbone amide or Gln11 and the carbonyl oxygen of the thioester group of malonyl-CoA, the distribution of distances between the backbone amide or Leu93 and the carbonyl oxygen of the thioester group of malonyl-CoA, the distribution of distances of Ser92's Oγ and the carbonyl carbon of thioester, the distribution of distances of Arg117, and the centers of geometry of α-carboxylate moiety of malonyl-CoA and Arg117's guanidinium function. The dotted vertical line indicates the average distance. Distances are measured in Å. 1D histograms was generated using a bin width of 0.25 Å.

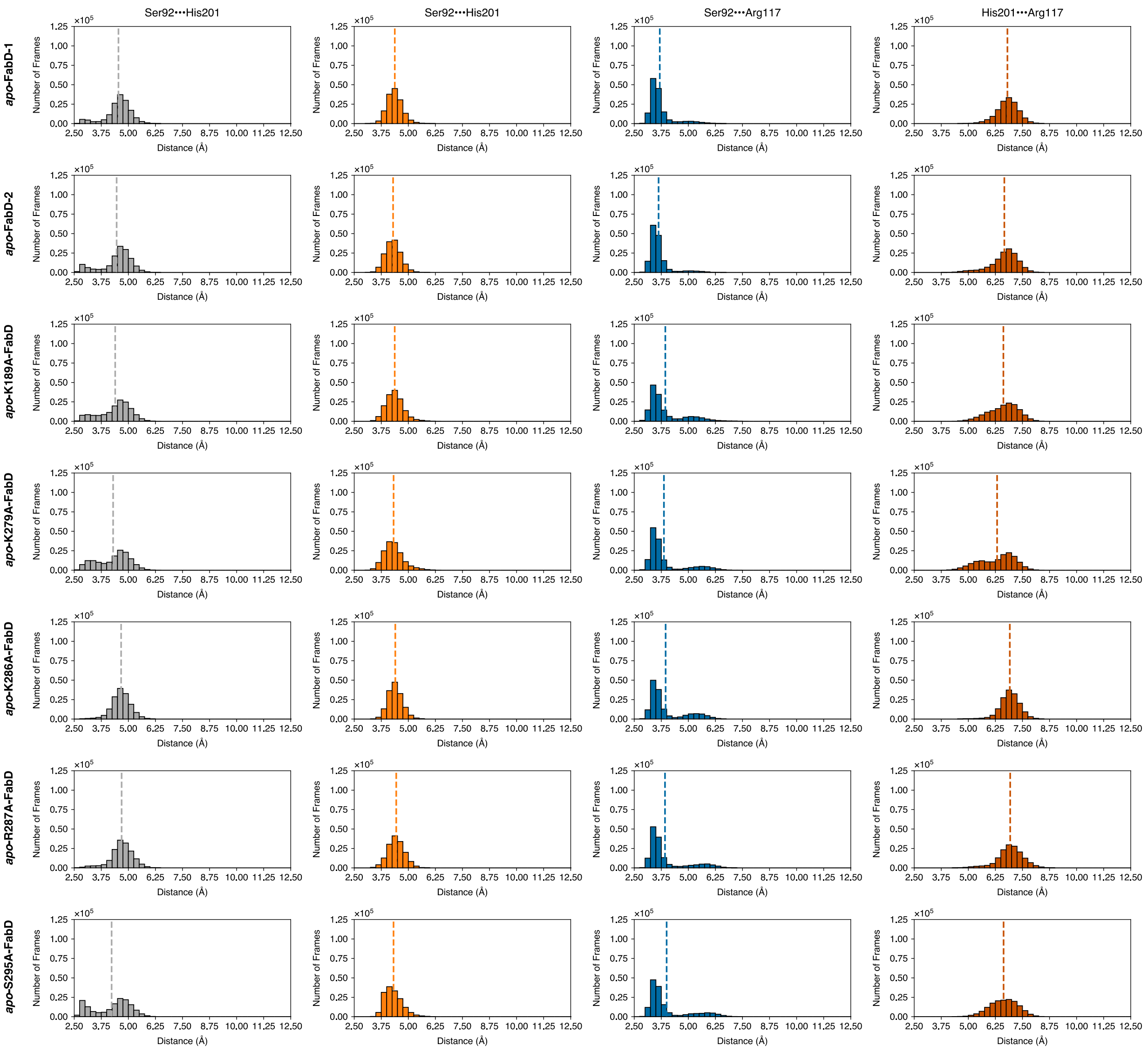

**Figure S25. Distribution of distances between key active site residues sampled throughout the course of MD simulations of *apo*-FabD variants.** From left to right the distribution of distances between Ser92 and His201 (measured in two different ways in the 1st and 2nd columns), Ser92 and Arg117, and His201 and Arg 117 (from left to right). The distances between Ser92 and His201 were measured as distance between Oγ of Ser92 and His92's Nε2 (1st column) or as the distance between Oγ of Ser and the center of geometry of His201's imidazole ring. The Ser92-Arg117 is measured between Oγ of Ser and the center of geometry of Arg117's guanidinium moiety. Lastly, the distance between His201 and Arg117 is determined the centers of geometry of the imidazole and guanidinium moiety of His201 and Argy117, respectively. The dotted vertical line indicates the average distance. Distances are measured in Å. 1D histograms was generated using a bin width of 0.25 Å.

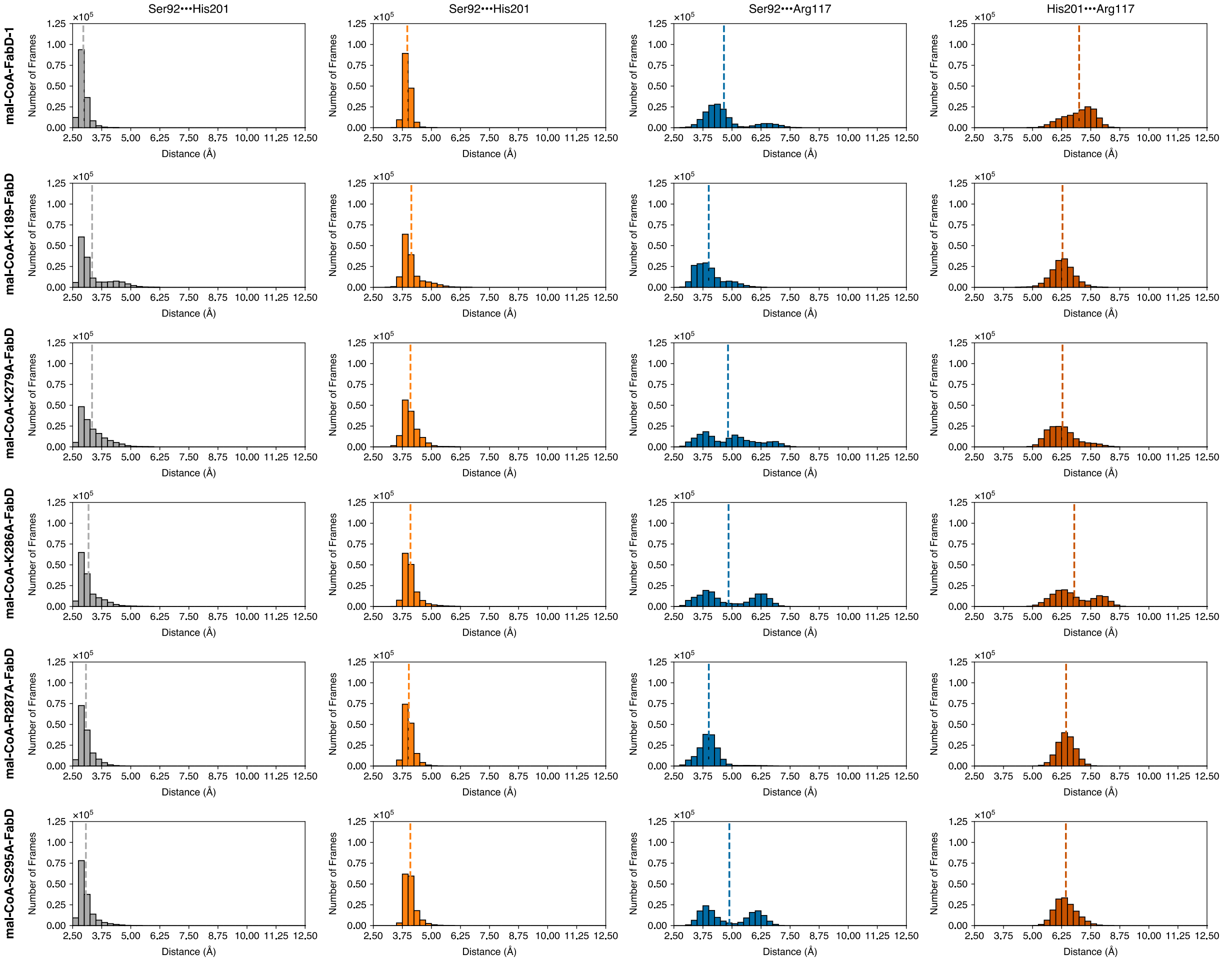

**Figure S26. Distribution of distances between key active site residues sampled throughout the course of MD simulations of malonyl-CoA·FabD variants.** From left to right the distribution of distances between Ser92 and His201 (measured in two different ways in the $1^{st}$ and $2^{nd}$ columns), Ser92 and Arg117, and His201 and Arg 117 (from left to right). The distances between Ser92 and His201 were measured as distance between Oγ of Ser92 and His92's Nε2 ($1^{st}$ column) or as the distance between Oγ of Ser and the center of geometry of His201's imidazole ring. The Ser92-Arg117 is measured between Oγ of Ser and the center of geometry of Arg117's guanidinium moiety. Lastly, the distance between His201 and Arg117 is determined the centers of geometry of the imidazole and guanidinium moiety of His201 and Argy117, respectively. The dotted vertical line indicates the average distance. Distances are measured in Å. 1D histograms was generated using a bin width of 0.25 Å.

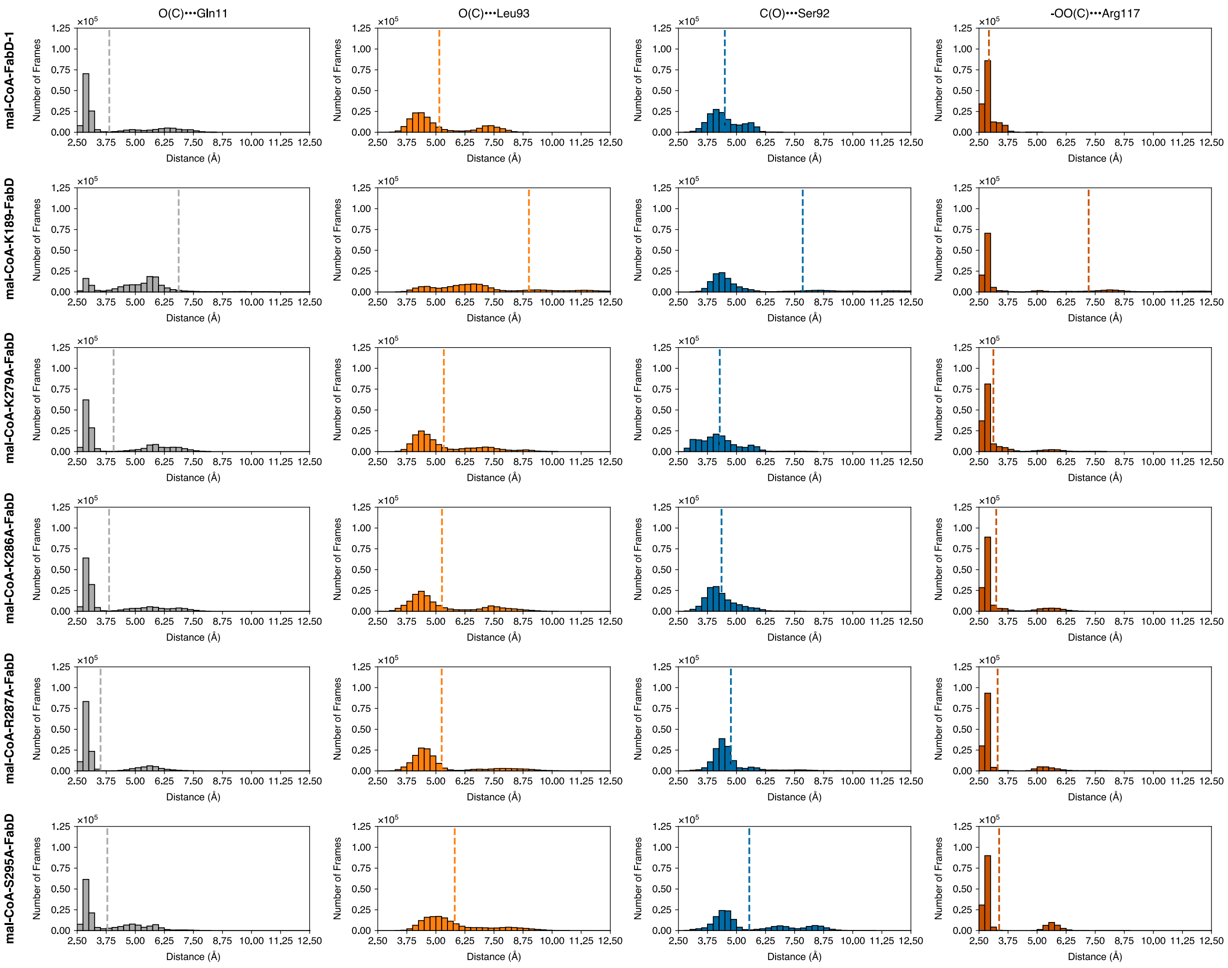

**Figure S27. Distribution of distances between key active site residues and malonyl portion of malonyl-AcpP sampled throughout the course of MD simulations of malonyl-CoA•FabD variants.** From left to right, the distribution of distances between the backbone amide or Gln11 and the carbonyl oxygen of the thioester group of malonyl-CoA, the distribution of distances between the backbone amide or Leu93 and the carbonyl oxygen of the thioester group of malonyl-CoA, the distribution of distances of Ser92's Oγ and the carbonyl carbon of thioester, the distribution of distances of Arg117, and the centers of geometry of α-carboxylate moiety of malonyl-CoA and Arg117's guanidinium function. The dotted vertical line indicates the average distance. Distances are measured in Å. 1D histograms was generated using a bin width of 0.25 Å.

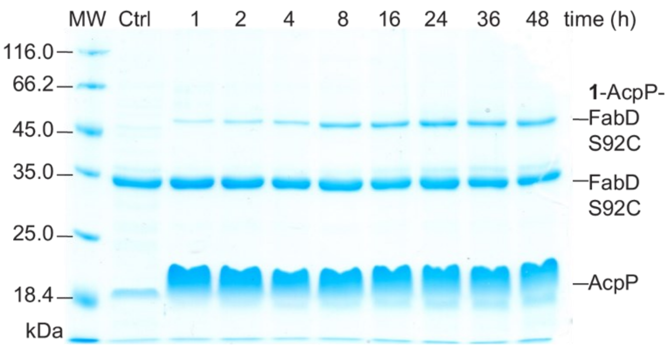

**Figure S28. Time-dependent C2α-bromo-*crypto*-AcpP-FabD S92C crosslinking.** SDS-PAGE analysis depicting crosslinking between 10 µM FabD S92C and 50 *µ*M C2-α-bromo-*crypto*-AcpP in PBS, pH 7 at 37 °C for 1–48 h. The lane denoted as Ctrl is a control reaction between 10 µM FabD and 20 µM *apo*-AcpP.

**Table S1. Table 1 X-ray crystallography data collection and refinement statistics**

| Primer name | Primer sequence |
|---|---|
| Complex | C2AcpP-FabD |
| Wavelength | 1 |
| Resolution range | 44.73 - 1.9 (1.968 - 1.9) |
| Space group | C 1 2 1 |
| Unit cell | 73.15 56.846 99.874 90 97.838 90 |
| Total reflections | 231636 (22001) |
| Unique reflections | 32206 (3205) |
| Multiplicity | 7.2 (6.9) |
| Completeness (%) | 99.87 (99.78) |
| Mean I/sigma(I) | 7.16 (1.53) |
| Wilson B-factor | 15.77 |
| R-merge | 0.286 (1.691) |
| R-meas | 0.335 (1.997) |
| R-pim | 0.173 (1.053) |
| CC1/2 | 0.987 (0.468) |
| CC* | 0.997 (0.805) |
| Reflections used in refinement | 32190 (3203) |
| Reflections used for R-free | 1704 (170) |
| R-work | 0.1846(0.3373) |
| R-free | 0.2213 (0.3647) |
| CC(work) | 0.959 (0.758) |
| CC(free) | 0.963 (0.712) |
| Number of non-hydrogen atoms | 3281 |
| macromolecules | 2874 |
| ligands | 57 |
| solvent | 350 |
| Protein residues | 385 |
| RMS(bonds) | 0.008 |

**Table S2. Molecular weight of FabD and *holo*-AcpP variants.** The molecular weight of each protein ($MW_{mea}$) was determined by LC ESI-TOF MS (Agilent 6230). The expected molecular weight of *holo*-AcpPs ($MW_{calc}$) have been calculated considering the $His_6$-tag cleavage and the installation of the PPant arm on AcpP's catalytic Ser36.

| **FabD** | $MW_{calc}$ (Da) | $MW_{mea}$ (Da) | **ΔMW (Da)** |
|---|---|---|---|
| WT | 34449.33 | 34449.00 | 0.33 |
| K189A | 34392.23 | 34391.70 | 0.53 |
| K189E | 34450.27 | 34449.90 | 0.37 |
| R190A | 34364.22 | 34364.20 | 0.02 |
| L192A | 34407.25 | 34407.20 | 0.05 |
| K279A | 34392.23 | 34392.20 | 0.03 |
| K279E | 34450.27 | 34450.00 | 0.27 |
| K286A | 34392.23 | 34391.80 | 0.43 |
| R287A | 34364.22 | 34364.00 | 0.22 |
| R287E | 34422.25 | 34421.80 | 0.45 |
| R287K | 34421.31 | 34421.20 | 0.11 |
| K189A R190A | 34307.12 | 34307.10 | 0.02 |
| K286A R287A | 34307.12 | 34306.50 | 0.62 |
| ***holo*-AcpP** | $MW_{calc}$ (Da) | $MW_{mea}$ (Da) | **ΔMW (Da)** |
| WT | 9392.33 | 9392.36 | 0.03 |
| E30A | 9334.30 | 9333.41 | 1.11 |
| D38A | 9348.32 | 9347.39 | 0.93 |
| E41A | 9334.30 | 9333.35 | 0.95 |

**Table S3. FabD systems subjected to molecular dynamics simulations**

| System | ff14SB/GAFF2 | CHARMM36m/CGenFF |
|---|---|---|
| *apo*-FabD-1 | Yes | Yes |
| *apo*-FabD-2 | Yes | Yes |
| *apo*-K189A-FabD | Yes | Yes |
| *apo*-K279A-FabD | Yes | Yes |
| *apo*-K286A-FabD | Yes | Yes |
| *apo*-R276A-FabD | Yes | Yes |
| *apo*-S295A-FabD | Yes | Yes |
| malonyl-CoA•FabD | Yes | Yes |
| malonyl-CoA-K189A-FabD | Yes | No |
| malonyl-CoA-K279A-FabD | Yes | No |
| malonyl-CoA-K286A-FabD | Yes | Yes |
| malonyl-CoA-R276A-FabD | Yes | Yes |
| malonyl-CoA-S295A-FabD* | Yes | Yes |
| malonyl-AcpP•FabD | Yes | Yes |
| malonyl-AcpP-K189A•FabD | Yes | Yes |
| malonyl-AcpP-K279A-FabD | Yes | Yes |
| malonyl-AcpP-K286A-FabD | Yes | Yes |
| malonyl-AcpP-R276A-FabD | Yes | Yes |
| malonyl-AcpP-S295A-FabD* | Yes | Yes |
| (malonyl-CoA•)FabD | Yes | Yes |
| (malonyl-AcpP•)FabD | Yes | Yes |

*S295 was a residue initially selected for mutation but ultimately not investigated experimentally. Nonetheless, MD simulations of the *apo*- and substrate-bound S295A-FabD systems were in progress and provide a considerable amount of additional data (2.286 µs). Because these data, in principle, improves sampling it was included in our computational analysis.

**Table S4. PCR primers used in this study.**

| Primer name | Primer sequence |
|---|---|
| LM_E30Af | 5’- CGT TGC GGA CCT GGG CGC GGA TTC -3’ |
| LM_E30Ar | 5’- GTC CGC AAC GAA AGA AGC ATT GTT GGT AAC TTC TTC CTG CTT AAC GCC -3’ |
| LM_E41Af | 5’- CGT TGC GCT GGT AAT GGC TCT GGA AGA AGA GTT TGA TAC TGA GAT TCC -3’ |
| LM_E41Ar | 5’- CAG CGC AAC GGT GTC AAG AGA ATC CGC GC -3’ |
| LM_R287Kf | 5’- CGA AAA AAA TTG TCG ACA CCC TGA CCG CCT CG -3’ |
| LM_R287Kr | 5’- CGA CAA TTT TTT TCG TCA GGC CAG TAA GCA CTT TGC C -3’ |
| LM_R287Af | 5’- CGA AAG CGA TTG TCG ACA CCC TGA CCG CC -3’ |
| LM_R287Ar | 5’- ACA ATC GCT TTC GTC AGG CCA GTA AGC ACT TTG CC -3’ |
| LM_R287Ef | 5’- CGA AAG AAA TTG TCG ACA CCC TGA CCG CCT CG -3’ |
| LM_R287Er | 5’- CGA CAA TTT CTT TCG TCA GGC CAG TAA GCA CTT TGC C -3’ |
| LM_K279Af | 5’- GGC GCG GTG CTT ACT GGC CTG ACG AAA C -3’ |
| LM_K279Ar | 5’- ACC GCG CCC GGG CCG ACT TCA TAG AGA TGT TC -3’ |
| LM_K279Ef | 5’- GGG CGA AGT GCT TAC TGG CCT GAC GAA ACG C -3’ |
| LM_K279Er | 5’- CAC TTC GCC CGG GCC GAC TTC ATA GAG ATG TTC TAC G -3’ |
| LM_K286Af | 5’- CTG ACG GCG CGC ATT GTC GAC ACC CTG -3’ |
| LM_K286Ar | 5’- GAC AAT GCG CGC CGT CAG GCC AGT AAG CAC -3’ |
| LM_L192Af | 5’- CGC GGC GCC GTT ACC AGT GAG CGT AC- 3’ |
| LM_L192Ar | 5’- GTA ACG GCG CCG CGC GTT TTG CG -3’ |
| LM_K189Af | 5’- GCA GCA CGC GCG CTG CCG TTA CCA G- 3’ |
| LM_K189Ar | 5’- GCG TGC TGC GCC CGC CGC TTT ACA G -3’ |
| LM_K189Ef | 5’- GCA GAA CGC GCG CTG CCG TTA CCA G- 3’ |
| LM_K189Er | 5’- GCG TTC TGC GCC CGC CGC TTT ACA G -3’ |
| LM_R190Af | 5’- GCA AAA GCG GCG CTG CCG TTA CCA GTG AGC -3’ |
| LM_R190Ar | 5’- GCC GCT TTT GCG CCC GCC GCT TTA CAG G -3’ |
| LM_K189AR190Af | 5’- GCA GCA GCA GCG CTG CCG TTA CCA GTG AG -3’ |
| LM_K189AR190Ar | 5’- GCT GCT GCT GCG CCC GCC GCT TTA C -3’ |
| LM_K286AR287Af | 5’- ACG GCG GCG ATT GTC GAC ACC CTG ACC -3’ |
| LM_K286AR287Ar | 5’- CGC CGC CGT CAG GCC AGT AAG CAC TTT G -3’ |

**Movie S1 (separate file).** A short video clip showing the pseudo-trajectory of the 1st principle component of FabD monomer of *apo*-FabD. Analysis was performed using all *apo*-FabD variants. The frames in which the ferredoxin-like (FL) domain is colored grey or black are the frames used in main text Figure 5.

**Movie S2 (separate file).** A short video clip showing the pseudo-trajectory of the 1st principle component of FabD monomer of malonyl-CoA•FabD. Analysis was performed using all malonyl-CoA•FabD variants. The frames in which the ferredoxin-like (FL) domain is colored grey or black are the frames used in main text Figure 5.

**Movie S3 (separate file).** A short video clip showing the pseudo-trajectory of the 1st principle component of FabD monomer of malonyl-AcpP•FabD. Analysis was performed using all malonyl-AcpP•FabD variants. The frames in which the ferredoxin-like (FL) domain is colored grey or black are the frames used in main text Figure 5.

### E. $^{1}H$ and $^{13}C$ NMR Spectra

1H-NMR (500 MHz) of PMB-protected pantetheineamide S2 in CDCl3

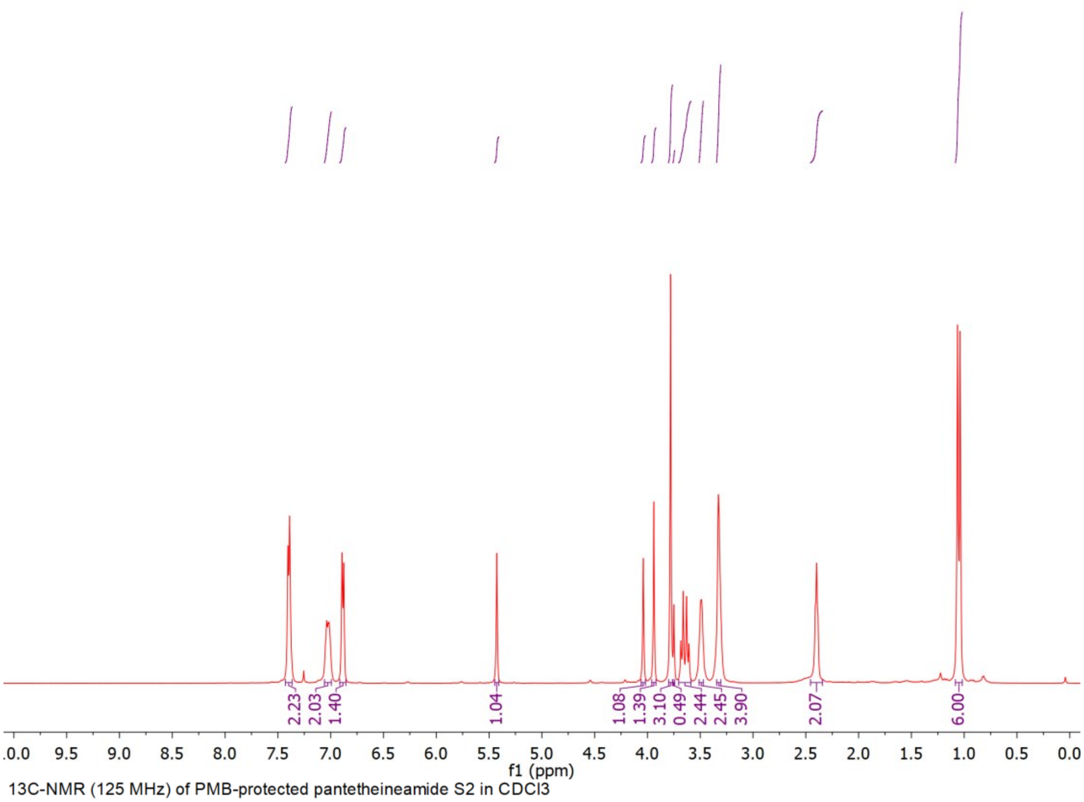

13C-NMR (125 MHz) of PMB-protected pantetheineamide S2 in CDCl3

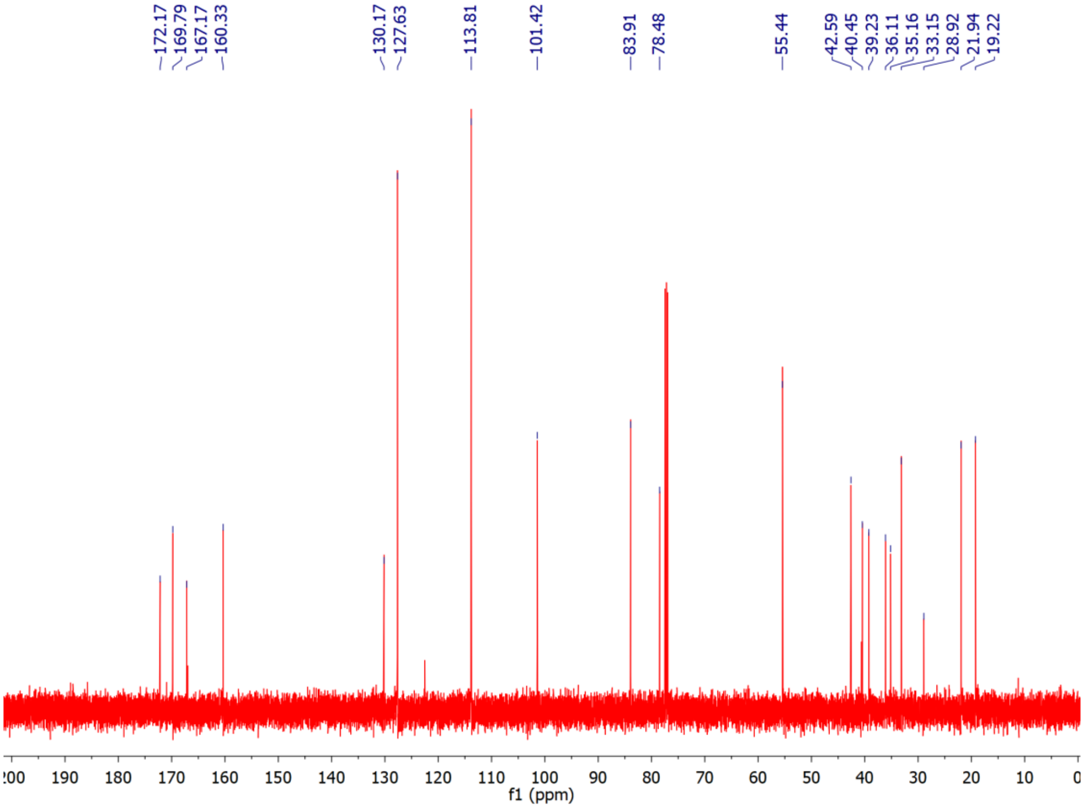

1H-NMR (400 MHz) of pantetheineamide 1 in CD3OD

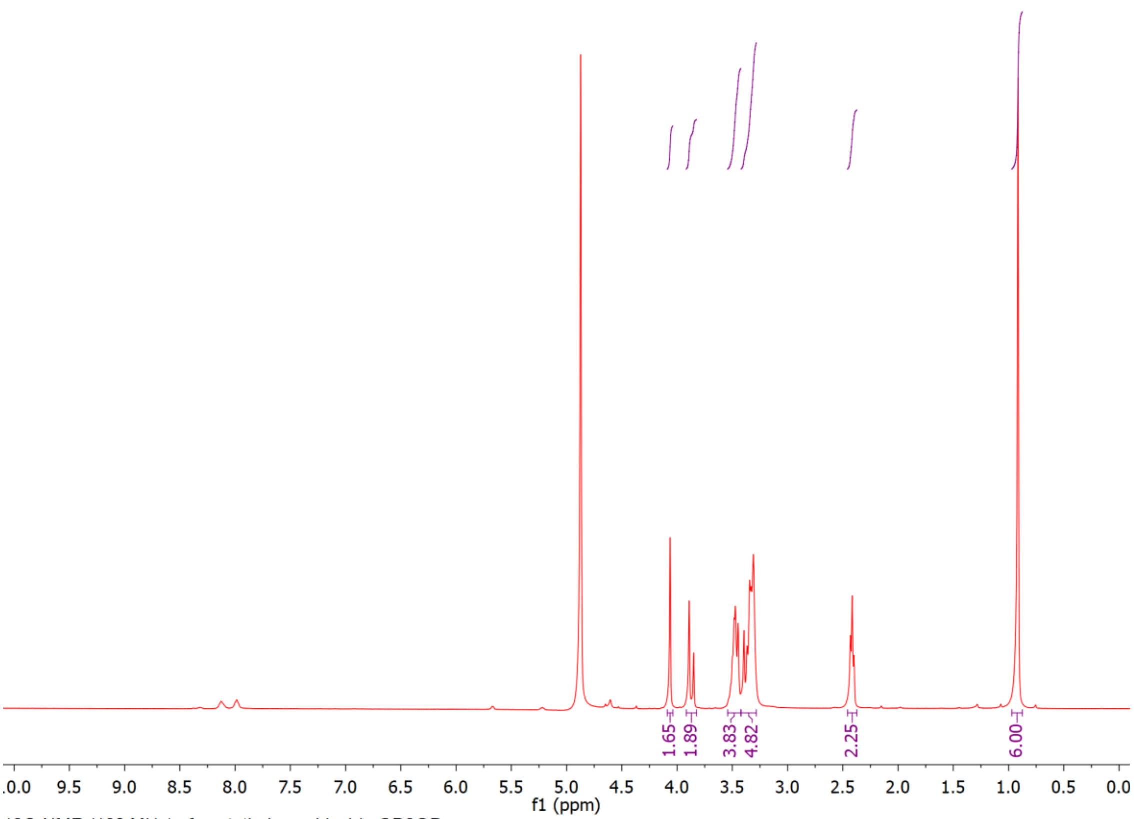

13C-NMR (100 MHz) of pantetheineamide 1 in CD3OD

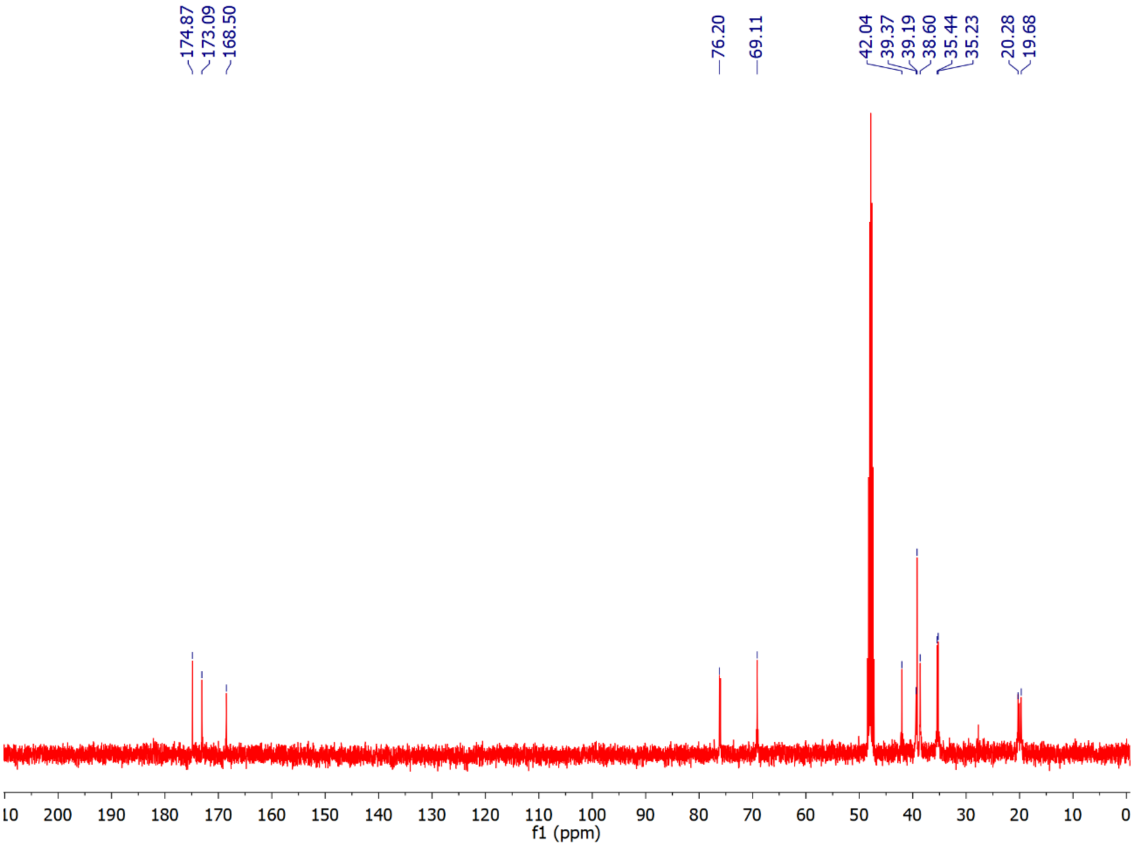
